# Cell type-selective AADC expression reduces motor deficits and dyskinesia in Parkinson’s disease mouse models

**DOI:** 10.64898/2026.01.20.700641

**Authors:** Ananya Chowdhury, Alex Fraser, Max Departee, Naz Taskin, Meagan A. Quinlan, John K. Mich, Victoria Omstead, Nathaly Lerma, Ximena Opitz-Araya, Alex C. Hughes, Emily Kussick, Refugio Martinez, Melissa Reding, Elizabeth Liang, Lyudmila Shulga, Dean Rette, Cindy Huang, Brittny Casian, Madison Leibly, Olivia Helback, Tyler Barcelli, Toren Wood, Nicholas Uribe, Colin Bacon, Jessica Bowlus, Dakota Newman, Rana Kutsal, Shannon Khem, Nicholas Donadio, Shenqin Yao, Kara Ronellenfitch, Vonn Wright, Kathryn Gudsnuk, Greg D. Horwitz, Boaz P. Levi, Ed S. Lein, Jonathan T. Ting, Tanya L. Daigle

**Affiliations:** Allen Institute for Brain Science, Seattle, WA 98109, USA; Washington National Primate Research Center, Seattle, WA 98195 USA; Department of Neurobiology and Biophysics, University of Washington, Seattle, WA 98195 USA

**Keywords:** AAV, AADC, neurodegeneration, Parkinson’s disease, L-DOPA, striatum, dopamine, nigrostriatal, motor behaviors, locus coeruleus, midbrain

## Abstract

Degeneration of midbrain dopamine (DA) neurons and the resulting loss of striatal dopamine signaling are hallmarks of Parkinson’s disease (PD). Although the dopamine precursor levodopa (L-DOPA) provides symptomatic relief, prolonged treatment often leads to abnormal involuntary movements (dyskinesia). Previous adeno-associated virus (AAV) approaches delivering aromatic L-amino acid decarboxylase (AADC) to the striatum under a ubiquitous promoter enhanced local dopamine synthesis and improved PD motor deficits, but this broad targeting strategy limited insight into the cellular populations underlying the behavioral improvements. Here, we engineered enhancer-driven AAVs to direct AADC expression to defined striatal and midbrain cell populations and paired these regulatory elements with a blood–brain barrier–penetrant capsid to enable both systemic and direct delivery. Targeted expression restored motor performance at reduced L-DOPA doses and decreased dyskinesia-like behaviors, producing improvements comparable to or greater than those achieved with ubiquitous expression. Distinct neuronal and non-neuronal populations each supported motor rescue but improved different behavioral domains to different extents, indicating complementary cell type-specific roles within PD-relevant circuits. Together, these findings establish enhancer-driven, cell type-specific AADC delivery as a better-tolerated strategy that enables rescue of motor deficits at lower L-DOPA doses in PD models.

## Introduction

Parkinson’s disease (PD) is a neurodegenerative disorder characterized by progressive degeneration of midbrain dopaminergic neurons and disruption of striatal dopamine (DA) signaling^1–3^, manifesting as severe motor impairments including rigidity, tremor, gait imbalance, grip difficulty, speech deficits, and micrographia^4,5^. Despite extensive understanding of the underlying cellular pathology, there is no cure or therapy that slows disease progression. The dopamine precursor levodopa (L-DOPA) remains the most widely used treatment for PD.

Although L-DOPA transiently restores dopamine signaling and improves motor function, prolonged treatment frequently leads to abnormal involuntary movements known as L-DOPA-induced dyskinesia (LID)^6–11^. These complications substantially reduce quality of life and highlight the need for strategies that restore dopamine signaling while minimizing adverse motor effects.

Advances in genetic engineering, viral vector development, and neurosurgical delivery have led to a rapid expansion of gene-based therapies for neurological disorders^12–16^. In dopamine-deficient conditions such as aromatic L-amino acid decarboxylase (AADC) deficiency and PD, promising results have been obtained using adeno-associated virus serotype 2 (AAV2) vectors to express AADC, a key enzyme required for dopamine synthesis^17,18^, under the ubiquitous cytomegalovirus (CMV) promoter^19–22^. The efficacy and safety of this AAV2-based gene therapy were demonstrated in children with AADC deficiency following delivery to both the putamen and midbrain^20–23^, leading to approval of this approach by the European Medicines Agency (EMA) and the U.S. Food and Drug Administration (FDA) for the treatment of pediatric AADC deficiency^24–26^. This strategy also increased striatal dopamine production, reduced L-DOPA requirements, and rescued motor deficits in rodents and non-human primates^27–31^. Clinical studies further showed that AAV2-AADC was generally well tolerated and improved motor function following bilateral intraputaminal infusions in adults with PD^32–35^.

Despite the therapeutic benefits of CMV-driven AADC strategies, the underlying cellular mechanisms remain unclear due to the ubiquitous nature of the promoter^36–38^. AADC expression in the largest striatal neuronal population, medium spiny neurons (MSNs), has been proposed as a primary mode of action^30,32,39^; however, intraputaminal delivery of CMV-driven AADC does not exclude contributions from other striatal cell types to the observed increases in extracellular dopamine (DA). In addition to MSNs, the striatum contains monoenzymatic nondopaminergic neurons that could synthesize and release DA following delivery of a functional DDC gene^40–42^. Ubiquitous AADC delivery can also transduce non-neuronal astrocytes, which play important roles in L-DOPA uptake and metabolism^43,44^ and can contribute to local DA production. DA generated in these transduced neuronal and non-neuronal populations may then diffuse through the extracellular space to activate dopamine receptors on neighboring striatal neurons or modulate intracellular signaling pathways, suggesting that multiple cell types could potentially contribute to the functional effects of AADC gene delivery.

Better definition of the roles of different cellular populations in affected brain regions may inform the development of more precise therapeutic strategies. Enhancer-based AAV technology is one approach that has been shown to be capable of driving robust, cell type-selective transgene expression to diverse neuronal and non-neuronal populations in the CNS using both direct and systemic delivery routes^45–50^. These tools have already been used successfully to correct severe circuit-level defects in genetic mouse models of epilepsy^51^, suggesting that enhancer-driven strategies could be applied across many disorders and translational contexts, including to target cell type- and disease-relevant circuits in PD.

Here, we generated enhancer-driven AAV vectors to direct AADC expression to selected striatal, midbrain, and non-neuronal cell populations. Characterization of these vectors in mice following direct and systemic delivery revealed distinct patterns of AADC expression across relevant brain regions. We then evaluated an MSN-targeted vector delivered in combination with low-dose L-DOPA and observed reversal of selected motor deficits in both 6-hydroxydopamine (6-OHDA) and MCI-Park PD mouse models using either route of administration. MSN-specific AADC expression produced phenotypic rescue comparable to or greater than that achieved with a ubiquitous promoter-driven vector and reduced L-DOPA–induced dyskinesia in the 6-OHDA model. Targeting striatal cholinergic neurons, midbrain dopaminergic neurons, and brain-wide astrocytes resulted in differential rescue of motor deficits in the rapid PD model (6-OHDA) but not in the progressive model (MCI-Park).

Together, these findings demonstrate that multiple neuronal and non-neuronal populations support motor recovery to different extents and with distinct roles, establishing enhancer-driven, cell type-specific AADC delivery as a strategy to reduce L-DOPA requirements and dyskinesia in PD mouse models.

## Results

### Development and validation of cell type-specific AADC vectors

To enable cell type-specific AADC expression in striatal MSNs, we utilized a previously characterized MSN enhancer (AiE0450h) shown to selectively express the fluorescent reporter SYFP2 in mouse striatum when placed upstream of a minimal β-globin promoter^48^ (AiE0450h-SYFP2; AiP12236; **Figure 1A**, left). SYFP2 was then replaced with a codon-optimized human *DDC* gene to generate the corresponding AADC expression vector (AiE0450h-AADC(no intron)-HA; AiP16390; **Figure 1—supplement figure 1A,** left).

**Figure 1:**
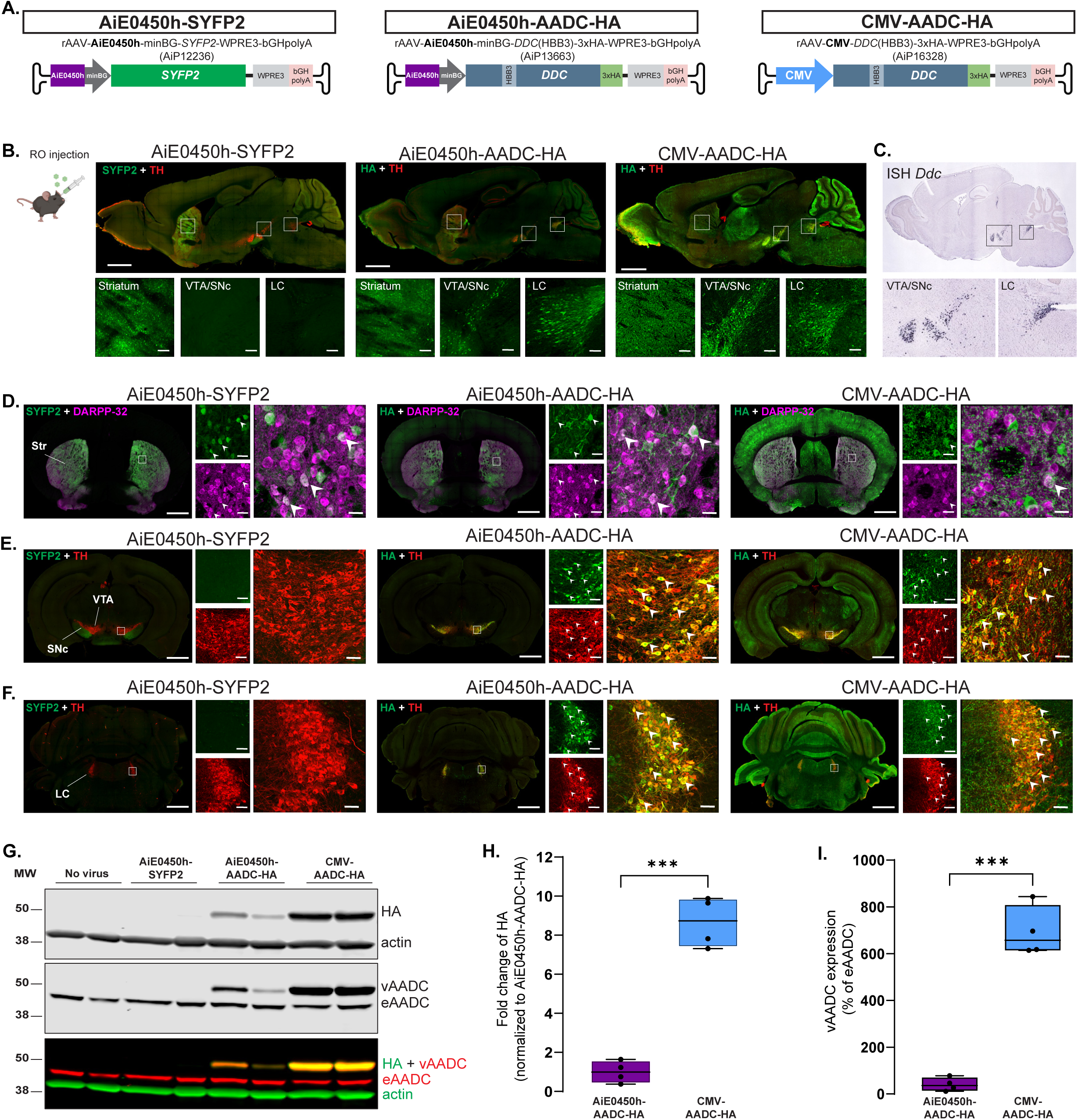
Development and validation of cell type-specific AADC vectors. (A) Schematic showing the vector designs for AiE0450h-SYFP2, AiE0450h-AADC-HA, and CMV-AADC-HA. (B) Schematic for retro-orbital (RO) injection and representative sagittal images of SYFP2 (green) and TH (red) for the enhancer control vector (AiE0450h-SYFP2; AiP12236, left); HA (green) and TH (red) for the cell type-specific vector (AiE0450h-AADC-HA; AiP13663, middle) and the ubiquitous promoter-driven vector (CMV-AADC-HA; AiP16328, right), following 1e12 vg RO administration. Top panels: low magnification (10X) stitched, scale bars, 500 µm; bottom panels: low magnification (10X) zoomed into the striatum, VTA/SNc, and LC in that order, scale bars, 50 µm. Box depicts ROI. TH co-staining was used to designate the relevant areas. (C) *Ddc* ISH data from the Allen Brain Atlas to visualize Ddc expression in VTA/SNc and LC. (D) Representative striatal (AP = +0.14 to +0.26 from bregma) coronal images of SYFP2 (green) and DARPP-32+ MSNs (magenta) for AiE0450h-SYFP2 (left); HA (green) and DARPP-32+ MSNs (magenta) for AiE0450h-AADC-HA (middle) and CMV-AADC-HA (right), following RO administration. (E) Representative midbrain (AP = -2.92 to -3.08 from bregma) and (F) LC (AP = -5.40 to -5.52 from bregma) coronal images of SYFP2 (green) and TH+ cells (red) for AiE0450h-SYFP2 (left); HA (green) and TH+ cells (red) for AiE0450h-AADC-HA (middle) and CMV-AADC-HA (right). Low magnification (10X) stitched, scale bars, 500 µm; High magnification (20X), scale bars, 10 µm for the striatum and 20 µm for VTA and LC. White arrows depict representative colocalized cells. (G) Representative western blot of striatal tissue punches from non-injected and injected mice showing HA and actin bands (top), AADC bands (middle), and merged image (bottom) with HA (green) and viral AADC (vAADC; red) overlaying. The endogenous AADC (eAADC) appears as a lower-molecular-weight band. Actin (green) was used as a loading control. (H) Quantification of HA expression normalized to AiE0450h-AADC-HA. Data represented as box plots indicating 25^th^ to 75^th^ percentiles and median as the solid line with whiskers denoting minimum and maximum values. n = 4 per condition. The *p* value were calculated by Welch’s t-test. ∗∗∗*p* < 0.001 (I) Quantification of vAADC expression normalized to eAADC. Data represented as box plots indicating 25th to 75th percentiles and median as the solid line with whiskers denoting minimum and maximum values. n = 4 per condition. The *p* value was calculated by Welch’s t-test. ∗∗∗*p* < 0.001

To optimize transgene expression, we generated three additional variants incorporating different introns: the human hemoglobin β third intron (HBB3;^51^AiE0450h-AADC(HBB3)-HA; AiP13663; **Figure 1A**, middle), the SV40 small-t intron^52^ (AiE0450h-AADC(SV40 small-t)-HA; AiP16391; **Figure 1—supplement figure 1A,** middle), and a longer SV40 intron^53^ (AiE0450h-AADC(SV40)-HA; AiP16392; **Figure 1—supplement figure 1A,** right). All constructs contained a C-terminal 3×HA epitope tag to enable immunodetection of virally expressed AADC.

To evaluate transgene expression in vivo, vectors were packaged in the blood-brain barrier (BBB)-penetrant PHP.eB capsid^54^ and delivered retro-orbitally (RO) to adult C57BL6/J mice (1e12 viral genomes (vg) per animal). Three weeks after injection, brains were collected and AADC expression in the striatum was assessed by anti-HA immunostaining.

The intronless construct and the SV40 small-t variant showed no detectable HA expression, while the longer SV40 intron supported only weak expression **(Figure 1—supplement figure 1B)**. In contrast, incorporation of the HBB3 intron produced robust transgene expression **(Figure 1B**, middle). Based on these results, the HBB3-containing construct (AiE0450h-AADC-HA) was selected for subsequent experiments.

We additionally generated a construct in which the same codon-optimized *DDC* sequence with the HBB3 intron was driven by the CMV promoter to directly compare ubiquitous AADC expression with cell type-specific vectors (CMV-AADC-HA; AiP16328; **Figure 1A**, right). We next compared the expression profiles of AiE0450h-SYFP2, AiE0450h-AADC-HA, and CMV-AADC-HA using immunohistochemistry. As previously reported, SYFP2 expression was restricted to MSNs in the striatum, with projections extending to the ventral tegmental area and substantia nigra (VTA/SN) (**Figure 1B**, left). The AiE0450h-AADC-HA vector resulted in robust striatal expression and detectable transgene-produced HA signal in the VTA/SNc and locus coeruleus (LC) (**Figure 1B**, middle), regions known to endogenously express *Ddc* according to in situ hybridization data from the Allen Brain Atlas (**Figure 1C**). Notably, these regions did not show cellular reporter expression with the AiE0450h-SYFP2 construct, suggesting that the extrastriatal signal observed with AiE0450h-AADC-HA is unlikely to be driven by enhancer activity, but may reflect properties of the AADC transgene itself. In contrast, CMV-AADC-HA drove widespread expression throughout the brain, including within the VTA/SNc and LC, consistent with the ubiquitous activity of the CMV promoter (**Figure 1B**, right).

To confirm MSN targeting, we co-stained brain sections for the HA epitope and the MSN marker DARPP-32^55^. AiE0450h-driven HA expression strongly overlapped with DARPP-32+ MSNs in the striatum, mirroring the pattern observed for AiE0450h-SYFP2 (white arrows, **Figure 1D**, left and middle). In contrast, CMV-driven HA expression was detected broadly across neuronal and non-neuronal populations, including but not limited to MSNs (**Figure 1D**, right).

Because HA signal was also observed in the VTA/SNc and LC, we next examined whether the transgene was expressed in catecholaminergic neurons by assessing co-localization with the marker tyrosine hydroxylase (TH)^56,57^. In contrast to the SYFP2 control (**Figure 1E-F**, left), both AiE0450h and CMV vectors showed robust co-labeling of transgene-produced HA with TH+ neurons in VTA, SNc, and LC (white arrows, **Figure 1E-F**, middle and right). Consistent with their weak expression in the striatum, the intronless construct produced no detectable HA signal in these regions (**Figure 1—supplement figure 1C-D,** left). In contrast, constructs containing the SV40 small-t intron or the longer SV40 intron produced some HA signal in these areas (**Figure 1—supplement figure 1C-D,** middle and right), indicating that the extrastriatal expression observed with the HBB3-containing vectors is unlikely to be driven by the intron itself.

To assess the broader biodistribution profiles of AiE0450h-SYFP2, AiE0450h-AADC-HA, and CMV-AADC-HA, we stained for HA in fixed sections from the spinal cord and selected peripheral tissues (heart, liver, spleen, and kidney) of RO-injected animals. HA signal in the spleen and kidney was at background level and comparable across no-virus control and all virus-treated groups. Sparse HA labeling was observed in the heart and liver following delivery of the AiE0450h-AADC-HA vector, with no detectable signal in the spinal cord relative to SYFP2 and no-virus controls. In contrast, the CMV-driven AADC vector produced robust labeling in the heart, liver, and spinal cord (**Figure 1—supplement figure 2A)**.

To quantitatively compare expression levels in the striatum, virally expressed HA-tagged AADC and total AADC protein were analyzed by Western blot. In both AiE0450h-AADC-HA and CMV-AADC-HA samples, HA and AADC signals were detected at the expected molecular weight, with the viral AADC band migrating slightly above endogenous AADC due to the C-terminal 3×HA tag (**Figure 1G**). Quantification revealed that the CMV promoter drove 8.66 ± 0.64-fold higher HA expression than the AiE0450h enhancer vector (**Figure 1H**). CMV-driven AADC expression exceeded physiological levels, reaching 693.0 ± 53.8% of endogenous protein (**Figure 1I**). In contrast, AiE0450h-AADC-HA produced AADC levels corresponding to 40.3 ± 14.3% of endogenous protein (**Figure 1I**). To assess off-target expression, we also analyzed a non-striatal region (M1 cortex). Both HA and AADC transgene bands were detected only in CMV-AADC-HA-injected animals, whereas no signal was detected in AiE0450h-AADC-HA or AiE0450h-SYFP2 animals **(Figure 1—supplement figure 2B**).

Together, these findings demonstrate that the AiE0450h enhancer-driven AADC vector maintains selective expression in striatal MSNs while also producing transgene expression in catecholaminergic neurons within endogenous *Ddc*-positive regions such as the VTA/SNc and LC following systemic administration. In contrast, the CMV-driven vector produces substantially higher and broadly distributed transgene expression throughout the brain and body, consistent with the known ubiquitous activity of the promoter.

### Retro-orbital delivery of AiE0450h-AADC rescues specific motor deficits in the 6-OHDA model

To assess the effects of RO delivery of cell type-specific AADC vectors on motor impairments, we used an established toxin mouse model of PD^58–60^. Adult C57BL/6J mice received a unilateral stereotaxic injection of 6-hydroxydopamine (6-OHDA) into the dorsal striatum to generate a rapid dopamine-depletion model. The retrogradely transported toxin caused severe degeneration of TH+ fibers at the injection site and substantial depletion of TH+ neurons in the injected hemisphere, with 85.55 ± 4.97% loss in the SNc and 45.37 ± 7.03% loss in the VTA relative to the contralateral hemisphere (**Figure 2—supplement figure 1A**).

**Figure 2:**
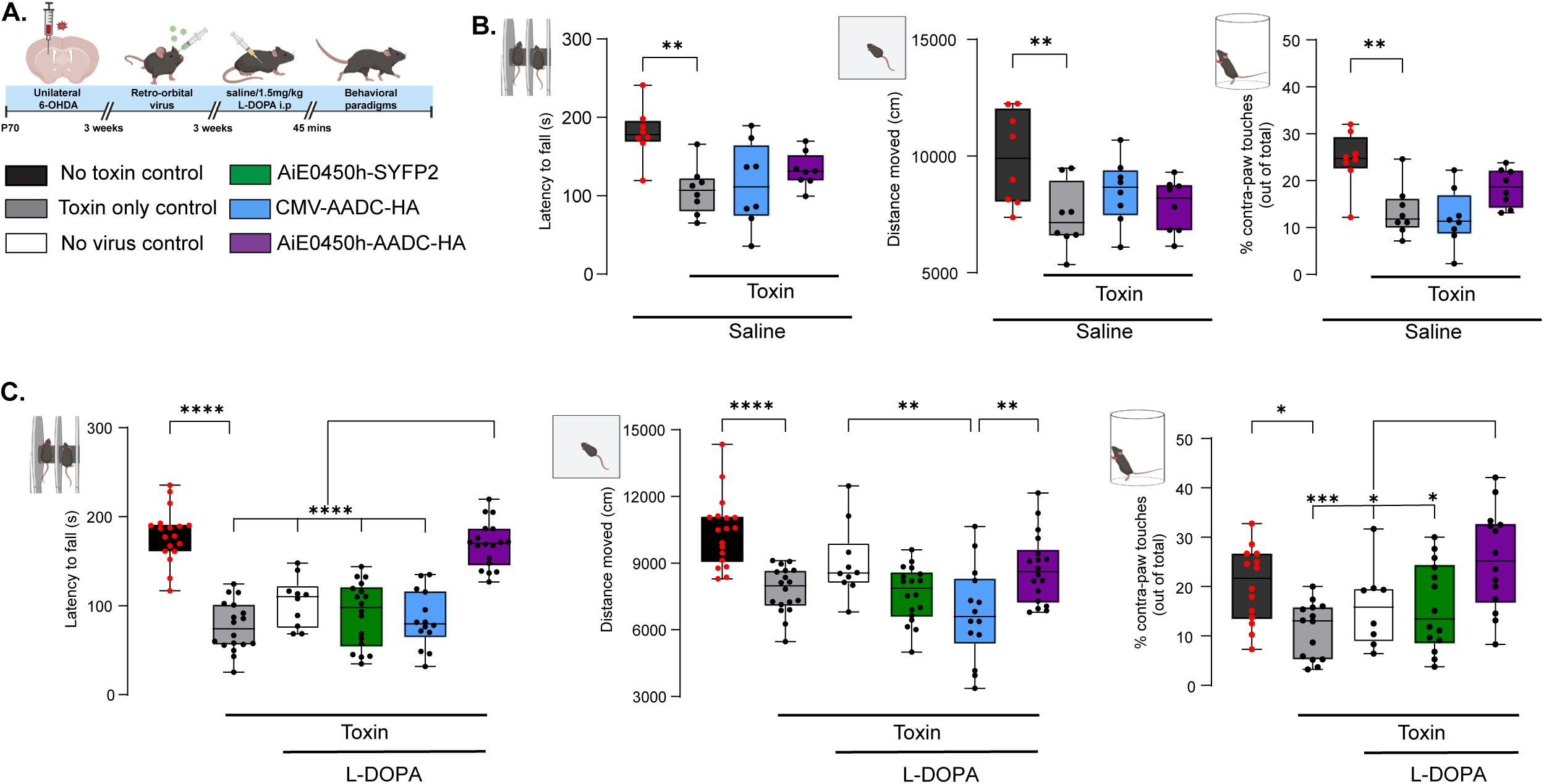
Retro-orbital delivery of AiE0450h-AADC rescues specific motor deficits in the 6-OHDA model. (A) Schematic overview of the injections and the behavioral tests. 6-OHDA toxin was stereotactically injected into the right dorsal striatum of P70 mice. Three weeks after the surgery, they received a RO injection of the viral vectors, and a series of behavioral tests were performed three weeks later, 45 mins after either saline or L-DOPA intraperitoneal injection as noted. (B) Retention time of the mice in the accelerating rotarod test (left), quantification of total distance traveled in the open field test (middle), and quantification of the first load-bearing use of contralateral paw (as percentage out of total paw touches) during rearing in a cylinder (right) in the absence of L-DOPA. Data represented as box plots indicating 25^th^ to 75^th^ percentiles and median as the solid line with whiskers denoting minimum and maximum values. n = 8 per group. The *p* values were calculated by one-way ANOVA followed by Dunnett’s post hoc test. ∗*p* < 0.05, ∗∗*p* < 0.01, ∗∗∗*p* < 0.001. (C) Retention time of the mice in the accelerating rotarod test (left), quantification of total distance traveled in the open field test (middle), and quantification of the first load-bearing use of contralateral paw (as percentage out of total paw touches) during rearing in a cylinder (right) in the presence of L-DOPA. Data represented as box plots indicating 25^th^ to 75^th^ percentiles and median as the solid line with whiskers denoting minimum and maximum values. n = 10-18 for rotarod and open field, n = 8-14 for paw touches. The *p* values were calculated by one-way ANOVA followed by Tukey’s post hoc test. ∗*p* < 0.05, ∗∗*p* < 0.01, ∗∗∗*p* < 0.001, ∗∗∗∗*p* < 0.0001.

Toxin-lesioned mice were then injected RO with vehicle, AiE0450h-SYFP2, CMV-AADC-HA, or AiE0450h-AADC-HA (1e12 vg per animal), and behavioral testing was performed three weeks later (**Figure 2A**). Motor behavior was evaluated using three assays: the accelerating rotarod to assess motor coordination, the open field test to measure exploratory behavior, and the cylinder test to examine forelimb asymmetry (**Figure 2B-C**).

Consistent with the lesion model, toxin-treated mice displayed significant motor deficits across all behavioral assays compared with non-lesioned controls (**Figure 2B**). Toxin treatment reduced latency to fall on the accelerating rotarod (no toxin control: 180.42 ± 11.97 s vs toxin-only control: 106.71 ± 11.06 s; p = 0.0012, left), decreased total distance traveled in the open field (9911.56 ± 709.57 cm vs 7413.06 ± 507.49 cm; p = 0.008, middle), and lowered the percentage of contralateral paw touches during supported rearing in the cylinder test (24.49 ± 2.12% vs 13.43 ± 1.91%; p = 0.001, right). In the absence of L-DOPA, RO delivery of AADC vectors alone did not improve these motor phenotypes. Compared with toxin-only controls, mice receiving CMV-AADC-HA or AiE0450h-AADC-HA showed no significant improvement in rotarod performance (CMV: 114.13 ± 18.90 s, AiE0450h: 132.79 ± 7.85 s), open-field activity (CMV: 8485.05 ± 498.16 cm, AiE0450h: 7868.74 ± 405.92 cm) or contralateral paw use (CMV: 12.03 ± 2.15%, AiE0450h: 18.51 ± 1.44%, **Figure 2B**).

Because AADC requires L-DOPA as a substrate, we next evaluated whether viral AADC expression could potentiate the effects of a subthreshold dose of L-DOPA. Toxin-lesioned mice were administered with a low dose of L-DOPA (1.5 mg/kg, intraperitoneal), and motor behavior was assessed 45 minutes later. At this dose, L-DOPA alone (no virus control) did not improve behavioral performance relative to toxin-only controls in the accelerating rotarod (toxin-only control: 76.41 ± 6.60 s vs no virus control: 104.07 ± 8.85 s, **Figure 2C**, left), open-field (7780.27 ± 243.31 cm vs 9036.22 ± 517.94 cm, **Figure 2C**, middle), or cylinder test (11.19 ± 1.52% vs 15.95 ± 2.92%, **Figure 2C**, right). In contrast, mice receiving AiE0450h-AADC-HA exhibited marked improvement in several behavioral measures following administration of low-dose L-DOPA. AiE0450h-AADC-HA–treated animals showed significantly increased latency to fall on the rotarod compared with all other virus-treated groups and controls (AiE0450h-AADC-HA: 168.82 ± 6.08 s; AiE0450h-SYFP2: 90.48 ± 8.49 s; CMV-AADC-HA: 84.52 ± 8.54 s; p < 0.0001; **Figure 2C**, left). These mice also displayed significantly higher contralateral forelimb use in the cylinder test relative to AiE0450h-SYFP2 mice (25.42 ± 2.71% vs 15.85 ± 2.37%; p = 0.026), toxin-only controls (p = 0.0002), and no virus controls (p = 0.0419; **Figure 2C**, right). Assessment of contralateral paw touches in the CMV-AADC-HA group was not possible because injected animals failed to rear.

In the open-field assay (**Figure 2C**, middle), AiE0450h-AADC-HA-treated animals did not significantly increase total distance traveled compared with AiE0450h-SYFP2 (AADC: 8734.77 ± 365.90 cm vs SYFP2: 7613.45 ± 292.24 cm) or the controls but performed significantly better than the CMV-AADC-HA group (6747.60 ± 566.09 cm, p = 0.0057).

At the matched titer of 1e12 vg per animal, toxin-lesioned mice receiving AiE0450h-AADC-HA showed lower mortality (12.5%) than those receiving CMV-AADC-HA (62.5%; **Figure 2—supplement figure 1B-C**). Furthermore, analysis of motor asymmetry due to dopaminergic imbalance between the lesioned and intact sides in the unilateral model^61,62^ revealed that AiE0450h-AADC-HA–treated animals exhibited no contralateral hindlimb dragging or rotation events during 10 minutes of observation, whereas CMV-AADC-HA-treated mice displayed frequent contralateral movements (12.50 ± 3.97 occurrences; 65.72 ± 26.56 s spent; p = 0.0046; **Figure 2—supplement figure 1D**). To assess the durability of AiE0450h-AADC-HA effect in the toxin model, behavioral testing was performed thirty-five weeks post–RO injection (P340), compared to the standard three-week time point (**Figure 2—supplement figure 1E**). Immunohistochemistry showed robust HA labeling in both the striatum and VTA/SNc, confirming the enduring expression of the transgene (**Figure 2—supplement figure 1F)**. At this extended time point, AiE0450h-AADC-HA-treated mice still showed improved rotarod performance (148.42 ± 10.75 s) relative to untreated toxin-lesioned mice (93.86 ± 7.76 s; p = 0.0036; **Figure 2—supplement figure 1G**, left). Consistent with observations at the three-week time point, open-field activity did not differ significantly between groups (11135.5 ± 834.70 cm vs 9393.24 ± 662.95 cm; p = 0.0939; **Figure 2—supplement figure 1G**, right). AiE0450h-AADC-HA-treated mice also exhibited higher body weight (36.38 ± 1.50 g) compared with untreated toxin-lesioned mice (32.14 ± 0.67 g, p=0.0306, data not shown), consistent with improved overall condition over time.

Together, these findings demonstrate that systemic delivery of cell type-specific AADC improves select motor phenotypes with low-dose L-DOPA and increases survival in the 6-OHDA model, compared with ubiquitous AADC expression under matched conditions, and produces durable behavioral effects over an extended period.

### Intrastriatal delivery of AiE0450h-AADC rescues specific motor deficits in the 6-OHDA model

To compare viral efficacy using a direct delivery approach, we injected 1e9 vg of AiE0450h-AADC-HA and CMV-AADC-HA bilaterally into the dorsal striatum. Three weeks post-injection, brains were harvested and processed for immunohistochemical analysis of viral transduction. Both vectors showed robust HA expression with comparable spread throughout the striatum (**Figure 3A**). Western blot analysis revealed higher HA levels in CMV-AADC-HA–treated mice, averaging 3.32 ± 1.35-fold greater expression than AiE0450h-AADC-HA (**Figure 3B-C**). Consistently, CMV-AADC-HA produced viral AADC expression corresponding to 1623 ± 683% above endogenous AADC, whereas AiE0450h-AADC-HA resulted in a 306 ± 54% increase **(Figure 3D**).

**Figure 3:**
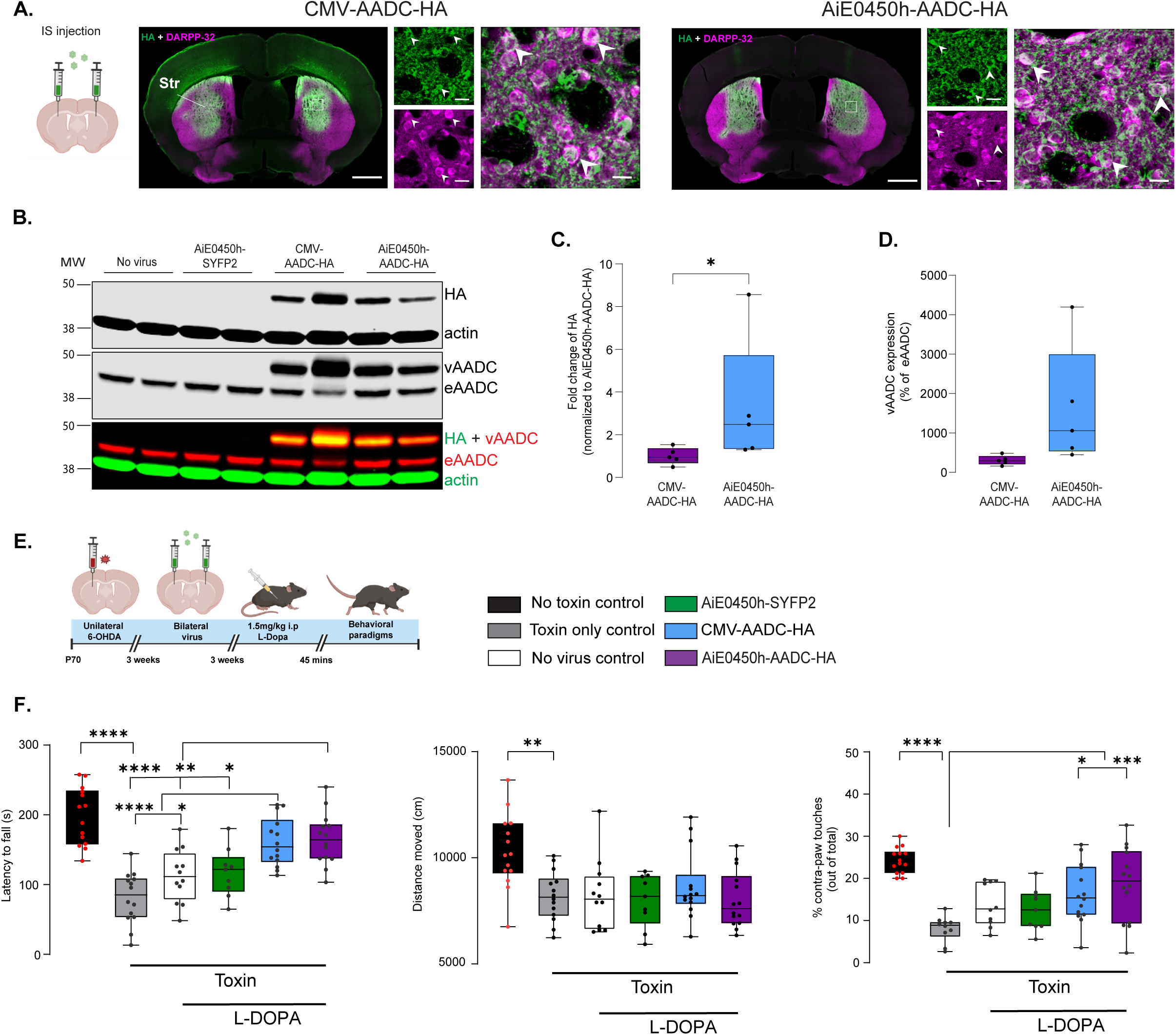
Intrastriatal delivery of AiE0450h-AADC rescues specific motor deficits in the 6-OHDA model. (A) Representative striatal (AP = +0.14 to +0.26 from bregma) coronal images of HA (green) and DARPP-32+ MSNs (magenta) for AiE0450h-AADC-HA (right panel) and CMV-AADC-HA (left panel), following 1e9 vg intrastriatal administration. Low magnification (10X) stitched, scale bars, 500 µm; High magnification (20X), scale bars, 10 µm. White arrows depict representative colocalized cells. (B) Representative western blot from striatal tissue punches showing expression of transgene produced HA tag and the loading control actin (top panel), viral AADC (vAADC) and endogenous AADC (eAADC) (middle panel). Bottom panel shows HA and vAADC overlap. (C) Quantification of fold-change in HA expression normalized to AiE0450h-AADC-HA. Data represented as box plots indicating 25^th^ to 75^th^ percentiles and median as the solid line with whiskers denoting minimum and maximum values. n = 5 per condition. The *p* value was calculated by Mann-Whitney test. ∗*p* < 0.05 (D) Quantification of vAADC expression normalized to eAADC. Data represented as box plots indicating 25th to 75th percentiles and median as the solid line with whiskers denoting minimum and maximum values. n = 5 per condition. The *p* value was calculated by Welch’s t-test. (E) Schematic overview of the injections and the behavioral tests. 6-OHDA toxin was stereotactically injected into the right dorsal striatum of P70 mice. Three weeks after the surgery, the viral vectors were administered bilaterally to the dorsal striatum, and a series of behavioral tests were performed three weeks later, 45 mins after L-DOPA intraperitoneal injection. (F) Retention time of the mice in the accelerating rotarod test (left), quantification of total distance traveled in the open field test (middle), and quantification of the first load-bearing use of contralateral paw (as percentage out of total paw touches) during rearing in a cylinder (right). Data represented as box plots indicating 25^th^ to 75^th^ percentiles and median as the solid line with whiskers denoting minimum and maximum values. n = 9-14 mice per condition. The *p* values were calculated by one-way ANOVA followed by Tukey’s post hoc test. ∗*p* < 0.05, ∗∗*p* < 0.01, ∗∗∗*p* < 0.001, ∗∗∗∗*p* < 0.0001.

At the injected dose of 1e9 vg per hemisphere, extrastriatal AADC expression was markedly reduced compared with previously shown systemic delivery (**Figure 3—supplement figure 1A-B**). Increasing the dose of AiE0450h-AADC-HA ten-fold per hemisphere increased extrastriatal HA expression, although levels remained substantially lower than those achieved with systemic delivery earlier (white arrows, **Figure 3—supplement figure 1C**). Together, these results indicate that direct injection produces robust AADC expression in striatal MSNs while substantially limiting extrastriatal expression.

To determine whether direct striatal delivery produced similar behavioral outcomes, we injected 1e9 vg of each vector bilaterally into the dorsal striatum of unilaterally toxin-lesioned mice and evaluated behavior three weeks later following administration of low-dose L-DOPA (1.5 mg/kg, i.p.; **Figure 3E**).

AiE0450h-AADC-HA-treated mice showed improved rotarod performance (165.17 ± 9.94 s), comparable to CMV-AADC-HA-treated animals (161.02 ± 9.38 s; p = 0.9997, **Figure 3F**, left). Both AADC vectors significantly increased their latency to fall compared to the toxin-only controls (81.05 ± 9.85 s; p < 0.0001) and no virus controls (112.61 ± 10.98 s, p=0.0083 for AiE0450h and p=0.0190 for CMV). AiE0450h-AADC-HA-treated mice also showed significant improvement in rotarod compared with AiE0450h-SYFP2 controls (117.33 ± 11.66 s; p = 0.0429), whereas CMV-AADC-HA-treated animals did not differ significantly from this control group (p = 0.0820).

Similarly, AiE0450h-AADC-HA-treated mice exhibited higher contralateral forelimb use in the cylinder test (18.61 ± 2.34%) comparable to CMV-AADC-HA-treated animals (16.36 ± 1.85%; p = 0.9121) and significantly greater than toxin-only controls (8.04 ± 0.97%, p= 0.0007 for AiE0450h and p= 0.0138 for CMV), but did not differ significantly from the no virus controls (13.74 ± 1.61%) and AiE0450h-SYFP2-treated mice (12.62 ± 1.67%, **Figure 3F**, right). In contrast, neither vector improved exploratory locomotion in the open-field assay (Toxin only control: 8188.89 ± 311.71 cm, no virus control: 8229.60 ± 476.50 cm, AiE0450h-SYFP2: 7994.12 ± 417.35 cm, CMV-AADC-HA: 8655.78 ± 427.24 cm, AiE0450h-AADC-HA: 8014.09 ± 356.57 cm, **Figure 3F**, middle). Importantly, intrastriatal delivery of either vector did not increase mortality or worsen behavioral phenotypes in toxin-lesioned mice (data not shown).

Together, these findings show that cell type-specific AADC improves motor performance in a PD model to a similar extent as ubiquitously expressed AADC following direct striatal delivery with low-dose L-DOPA. At the titers used, the enhancer-driven AAV largely restricted AADC expression to striatal MSNs, indicating that MSN-targeted AADC expression alone may be sufficient to drive motor rescue.

### AiE0450h-AADC reduces L-DOPA-induced dyskinesia in 6-OHDA model

Because long-term L-DOPA administration frequently produces dyskinesia in PD patients and animal models^6–11^, we next examined whether cell type-specific AADC expression affects L-DOPA-induced dyskinesia (LID).

To identify an appropriate L-DOPA dose for studying LID in toxin-lesioned mice, we first performed a dose-behavior response analysis using a single intraperitoneal injection of L-DOPA (**Figure 4A**). Relative to saline (0.00 ± 8.90%), doses of 3 mg/kg (48.76 ± 10.46%; p = 0.0174), 5 mg/kg (61.64 ± 7.76%; p = 0.0017), and 10 mg/kg (36.44 ± 12.68%; p = 0.0490) significantly improved rotarod performance in the toxin-lesioned mice (**Figure 4B**, left). Similarly, open-field activity increased at 3 mg/kg (25.82 ± 7.31%; p = 0.0339) and 5 mg/kg (29.47 ± 7.03%; p = 0.0119) relative to saline controls (0.0 ± 4.79%) (**Figure 4B**, middle). In contrast, both 1.5 mg/kg (low) and 20 mg/kg (high) doses failed to improve motor performance (**Figure 4B**).

**Figure 4:**
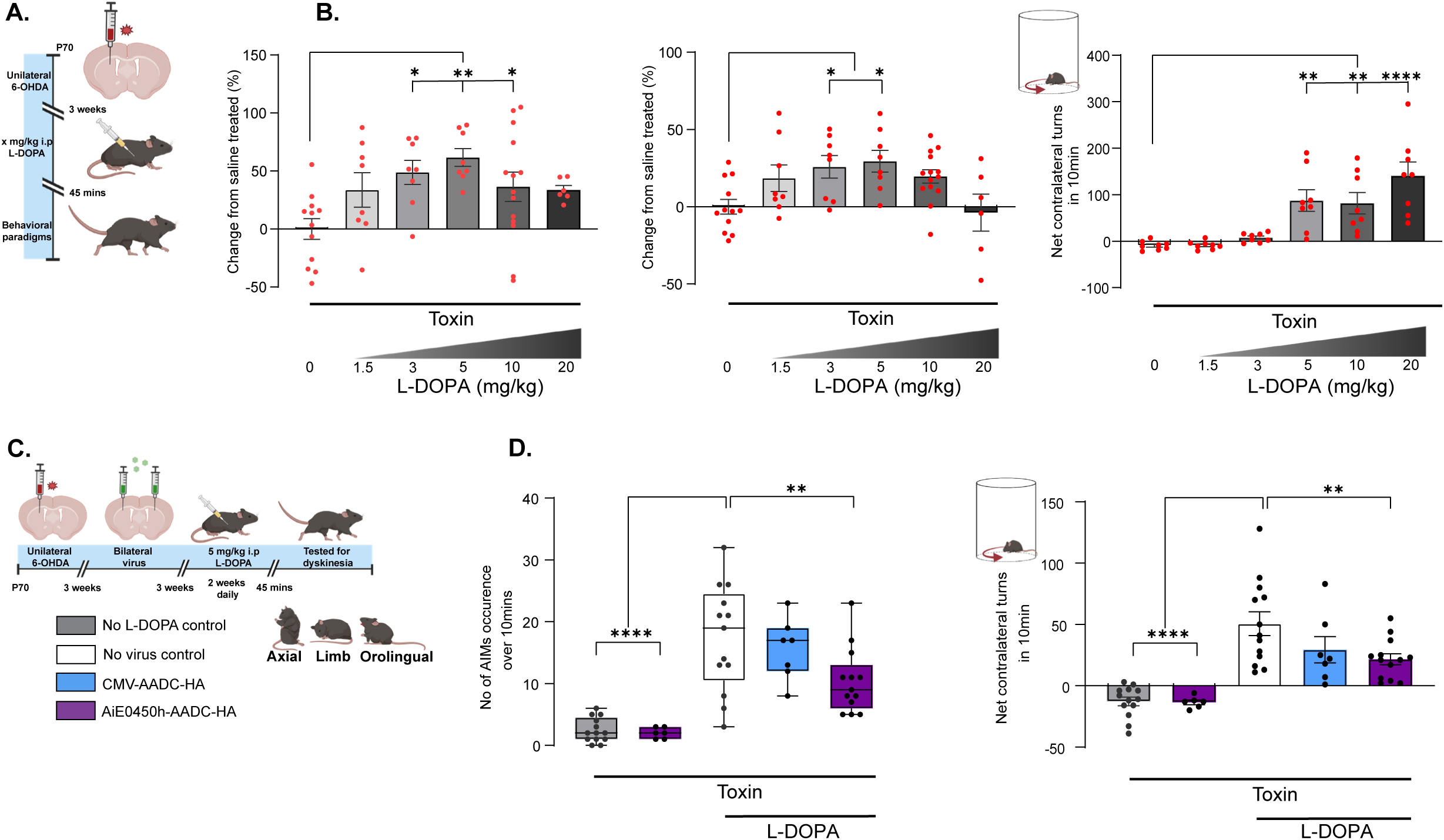
AiE0450h-AADC reduces L-DOPA-induced dyskinesia in 6-OHDA model. (A) Schematic overview of the injections and the behavioral tests. 6-OHDA toxin was stereotactically injected into the right dorsal striatum of P70 mice. 3 weeks after the surgery, a series of behavioral tests were performed 45 mins after either saline or different doses of L-DOPA intraperitoneal injection as noted. (B) Retention time of the mice in the accelerating rotarod test (left) and quantification of total distance traveled in the open field test (middle) (as percentage change from saline-injected, toxin-treated mice). Quantification of the net contralateral rotations in a cylinder over 10 mins (right). Data represented as mean ± SEM. n = 6-14 mice per condition. The *p* values were calculated by one-way ANOVA followed by Dunnett’s post hoc test. ∗*p* < 0.05, ∗∗*p* < 0.01, ∗∗∗∗*p* < 0.0001. (C) Schematic overview of the injections and the behavioral tests. 6-OHDA toxin was stereotactically injected into the right dorsal striatum of P70 mice. 3 weeks after the surgery, the viral vectors were administered bilaterally to the dorsal striatum, and intraperitoneal L-DOPA was administered daily for 2 weeks before testing dyskinesia (axial, limb, and orolingual). (D) Quantification of L-DOPA induced dyskinesia (LID) as the cumulation occurrence of abnormal involuntary movements (AIMs) affecting axial, limb, and orolingual muscles (left) and net contralateral rotations (right) in a cylinder over 10 mins. Data represented as box plots indicating 25^th^ to 75^th^ percentiles and median as the solid line with whiskers denoting minimum and maximum values and as mean ± SEM for rotations. N = 6-13 mice per condition. The *p* values were calculated by one-way ANOVA followed by Dunnett’s post hoc test. ∗∗*p* < 0.01, ∗∗∗∗*p* < 0.0001.

Consistent with the unilateral lesion model, higher L-DOPA doses also produced increasing contralateral rotations, known to be caused by dopamine receptor hypersensitivity on the lesioned side^63^. Significant rotational behavior occurred at 5 mg/kg (87.38 ± 23.17; p = 0.0023), 10 mg/kg (81.50 ± 22.93; p = 0.0044), and 20 mg/kg (140.75 ± 29.63; p < 0.0001), whereas lower doses did not differ in contralateral turning (1.5 mg/kg: −9.00 ± 3.06; 3 mg/kg: 7.50 ± 3.46) relative to saline controls (−9.38 ± 3.38) (**Figure 4B**, right).

Because 5 mg/kg L-DOPA produced strong behavioral improvement while inducing contralateral rotations, we used this dose to examine the effects of prolonged L-DOPA exposure in the presence of virally delivered AADC. To directly compare AiE0450h-AADC-HA and CMV-AADC-HA within the striatum, we delivered the vectors by bilateral intrastriatal injection. Three weeks after unilateral 6-OHDA lesioning, mice received 1e9 vg per hemisphere of each vector, followed by daily L-DOPA (5 mg/kg, i.p.) for two weeks (**Figure 4C**). On the final day, dyskinetic abnormal involuntary movements (AIMs) affecting axial, limb, and orolingual (ALO) muscles were scored for 10 minutes in the cylinder 45 minutes after L-DOPA administration.

Mice receiving L-DOPA for two weeks without viral treatment exhibited significantly higher cumulative AIM scores (17.31 ± 2.40) compared with saline-treated controls without virus (2.46 ± 0.55; p < 0.0001) or saline-treated mice receiving AiE0450h-AADC-HA (2.00 ± 0.37; p < 0.0001). When treated with the same prolonged L-DOPA regimen, mice that received AiE0450h-AADC-HA showed significantly reduced AIM scores (10.38 ± 1.47; p = 0.0093) compared with L-DOPA without viral treatment. In contrast, CMV-AADC-HA did not significantly reduce AIM scores (15.57 ± 1.88; p = 0.9162), although the difference between the two viral groups was not significant (p = 0.281, **Figure 4D**, left).

The same prolonged L-DOPA regimen also increased contralateral rotations in no virus group (50.62 ± 9.73; p < 0.0001) relative to saline controls (no virus: −12.77 ± 3.54, AiE0450h-AADC-HA: -13.50 ± 1.95). Bilateral intrastriatal administration of AiE0450h-AADC-HA significantly reduced this L-DOPA-induced rotational behavior (21.62 ± 4.54; p = 0.0088), whereas CMV-AADC-HA had no significant effect (29.43 ± 10.70; p = 0.1776) (**Figure 4D**, right).

Together, these results show that cell type-specific AADC expression reduces dyskinesia-like behaviors associated with prolonged L-DOPA exposure in the 6-OHDA model.

### AiE0450h-AADC improves motor phenotypes in the MCI-Park model

To determine whether selective viral AADC expression could influence motor phenotypes in a progressive PD model, we used a genetic mouse model in which the gene encoding an essential subunit of mitochondrial complex I, Ndufs2, is selectively deleted in dopaminergic neurons^64^. Floxed *Ndufs2* mice (*Ndufs2*^fl/fl^) were crossed with mice expressing Cre recombinase under the dopamine transporter promoter (DAT, encoded by *Slc6a3*; *Slc6a3*^+/cre^) to generate conditional knockouts (*Slc6a3*^+/cre^; *Ndufs2*^fl/fl^; hereafter referred to as MCI-Park mice). Littermates lacking Cre recombinase (*Slc6a3*^+/+^; *Ndufs2*^fl/fl^) were used as controls. Consistent with previous reports, MCI-Park mice showed pronounced bilateral loss of TH+ projections in the dorsal striatum and dopaminergic neuron loss in the VTA/SNc by P60 (**Figure 5—supplement figure 1A**). Because degeneration of TH+ axons is already detectable from P30^64^, mice were injected RO at P30-35 with vehicle, AiE0450h-SYFP2, CMV-AADC-HA, or AiE0450h-AADC-HA (1e12 vg) and evaluated for their behavioral performance four weeks later (P58-63) (**Figure 5A**).

**Figure 5:**
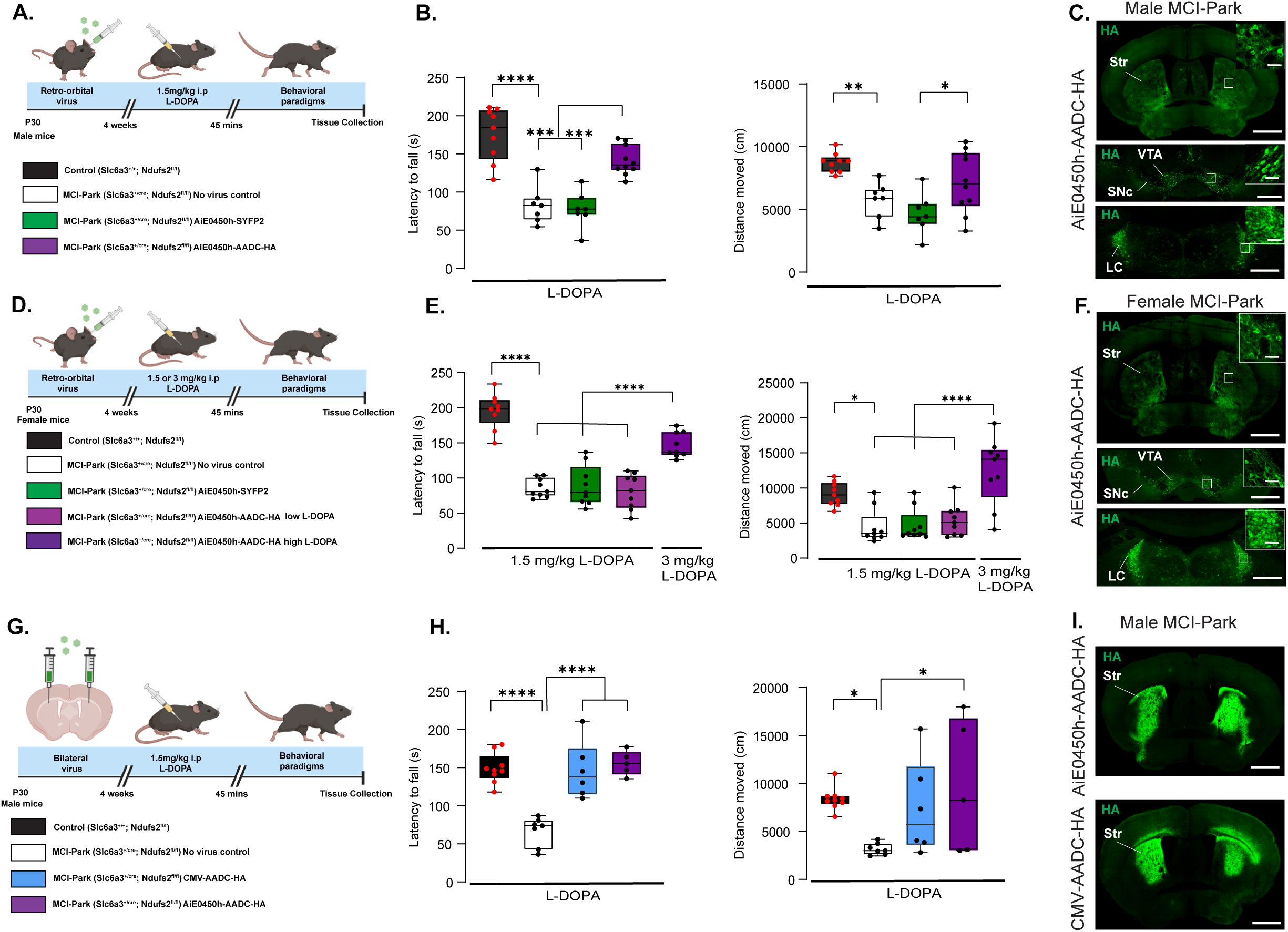
AiE0450h-AADC improves motor phenotypes in the MCI-Park model. (A) Schematic overview of the injections and the behavioral tests. Male MCI-Park (*Slc6a3*^+/cre^; *Ndufs2*^fl/fl^) and age-matched littermate controls (*Slc6a3*^+/+^; *Ndufs2*^fl/fl^) were RO injected with either the viral vectors or the vehicle at P30-35. 4 weeks later at around P60, a series of behavioral tests were performed 45 mins after L-DOPA intraperitoneal injection. (B) Retention time of the male mice in the accelerating rotarod test (left) and quantification of total distance traveled in the open field test (right). Data represented as box plots indicating 25^th^ to 75^th^ percentiles and median as the solid line with whiskers denoting minimum and maximum values. n = 7-10 mice per condition. The *p* values were calculated by one-way ANOVA followed by Tukey’s post hoc test. ∗*p* < 0.05, ∗∗*p* < 0.01, ∗∗∗*p* < 0.001, ∗∗∗∗*p* < 0.0001. (C) Representative coronal images of striatum (AP = +0.26 from bregma, top), midbrain (AP = -3.16 from bregma, middle) and LC (AP = -5.40 from bregma, bottom) showing HA (green) following RO administration of AiE0450h-AADC-HA. Low magnification (10X) stitched, scale bars, 500 µm, top; High magnification (20X), scale bars, 20 µm as insets. (D) Schematic overview of the injections and the behavioral tests. Female MCI-Park (*Slc6a3*^+/cre^; *Ndufs2*^fl/fl^) and age-matched littermate controls (*Slc6a3*^+/+^; *Ndufs2*^fl/fl^) were RO injected with either the viral vectors or the vehicle at P30-35. 4 weeks later at around P60, a series of behavioral tests were performed 45 mins after L-DOPA intraperitoneal injection at the doses mentioned. (E) Retention time of the female mice in the accelerating rotarod test (left) and quantification of total distance traveled in the open field test (right). Data represented as box plots indicating 25^th^ to 75^th^ percentiles and median as the solid line with whiskers denoting minimum and maximum values. n = 9 mice per condition. The *p* values were calculated by one-way ANOVA followed by Tukey’s post hoc test. ∗*p* < 0.05, ∗∗*p* < 0.01, ∗∗∗*p* < 0.001, ∗∗∗∗*p* < 0.0001. (F) Representative coronal images of striatum (AP = +0.26 from bregma, top), midbrain (AP= -3.16 from bregma, middle) and LC (AP= -5.40 from bregma, bottom) showing HA (green) following RO administration of AiE0450h-AADC-HA. Low magnification (10X) stitched, scale bars, 500 µm, top; High magnification (20X), scale bars, 20 µm as insets. (G) Schematic overview of the injections and the behavioral tests. The viral vectors or the vehicle was bilaterally injected into the dorsal striatum of male MCI-Park (*Slc6a3*^+/cre^; *Ndufs2*^fl/fl^) and age-matched littermate controls (*Slc6a3*^+/+^; *Ndufs2*^fl/fl^) at P30-35. 4 weeks later at around P60, a series of behavioral tests were performed 45 mins after L-DOPA intraperitoneal injection. (H) Retention time of the mice in the accelerating rotarod test (left) and quantification of total distance traveled in the open field test (right). Data represented as box plots indicating 25^th^ to 75^th^ percentiles and median as the solid line with whiskers denoting minimum and maximum values. n = 5-9 mice per condition. The *p* values were calculated by one-way ANOVA followed by Dunnett’s post hoc test. ∗*p* < 0.05, ∗∗∗∗*p* < 0.0001. (I) Representative coronal images of striatum (AP = +0.14 from bregma) for AiE0450h-AADC-HA (top) and CMV-AADC-HA (bottom) showing HA (green). Low magnification (10X) stitched, scale bars, 500 µm.

At the previously established low dose of L-DOPA (1.5 mg/kg), male MCI-Park mice without viral treatment showed significant motor deficits compared with age-matched controls, including reduced latency to fall on the accelerating rotarod (82.57 ± 9.25 s vs 175.74 ± 11.60 s; p < 0.0001, **Figure 5B**, left) and decreased distance traveled in the open field (5759.19 ± 530.03 cm vs 8717.00 ± 256.82 cm; p = 0.0100) (**Figure 5B**, right).

Male MCI-Park mice receiving AiE0450h-AADC-HA showed significant improvement in rotarod performance (141.27 ± 6.15 s; p = 0.0005) compared with no virus controls and AiE0450h-SYFP2–treated mice (77.24 ± 8.93 s; p = 0.0002) (**Figure 5B**, left). These mice also showed increased exploratory activity in the open-field assay (7166.63 ± 774.61 cm; p = 0.0308) compared with AiE0450h-SYFP2 controls (4671.06 ± 609.93 cm, **Figure 5B**, right). Systemic delivery of CMV-AADC-HA at the same 1e12 vg dose caused high mortality in MCI-Park mice, preventing reliable behavioral assessment (**Figure 5—supplement figure 1B**). Robust HA expression was detected in the striatum, VTA/SNc, and LC of male MCI-Park mice receiving AiE0450h-AADC-HA (**Figure 5C**).

Female MCI-Park mice (**Figure 5D**) without viral treatment similarly displayed motor deficits compared with controls, including reduced rotarod performance (86.04 ± 4.44 s vs 195.59 ± 8.33 s; p < 0.0001) and decreased open-field activity (4462.48 ± 808.74 cm vs 9130.89 ± 563.27 cm; p = 0.0121) (**Figure 5E**). However, in contrast to males, AiE0450h-AADC-HA did not improve motor performance in females at 1.5 mg/kg L-DOPA. Increasing the L-DOPA dose to 3 mg/kg significantly improved rotarod performance (146.56 ± 5.91 s; p < 0.0001) relative to no virus mice, AiE0450h-SYFP2 controls (88.96 ± 9.54 s), and AiE0450h-AADC-HA–treated mice at the lower L-DOPA dose (80.70 ± 8.26 s) (**Figure 5E**, left). At this dose, AiE0450h-AADC-HA–treated females also showed significantly increased open-field activity (12428.50 ± 1594.72 cm; p < 0.0001) compared with the no virus group and the mice that received AiE0450h-SYFP2 (4690.67 ± 764.72 cm) and AiE0450h-AADC-HA at low-dose L-DOPA (5390.75 ± 756.30 cm, **Figure 5E**, right). As observed in males, systemic CMV-AADC-HA delivery also increased mortality in female mice at the matched 1e12 vg dose (data combined in **Figure 5—supplement figure 1B**). Robust HA expression was detected in all three areas of female MCI-Park mice receiving AiE0450h-AADC-HA (**Figure 5F**).

To compare viral vectors using direct delivery, bilateral intrastriatal injections (1e9 vg per hemisphere) were performed in MCI-Park male mice at P30-35, followed by behavioral testing four weeks later using 1.5 mg/kg L-DOPA (**Figure 5G**).

AiE0450h-AADC-HA–treated mice showed rotarod performance (155.67 ± 7.14 s) comparable to CMV-AADC-HA–treated mice (146.01 ± 15.17 s; p = 0.9096) and both groups performed significantly better than MCI-Park mice without viral treatment (66.52 ± 7.30 s; p < 0.0001) (**Figure 5H**, left). AiE0450h-AADC-HA–treated mice also showed significantly greater open-field activity (9581.95 ± 3111.27 cm; p = 0.0383) compared with no virus group (3167.62 ± 235.42 cm), whereas CMV-AADC-HA did not produce a significant increase (7362.13 ± 2024.14 cm; p = 0.2206, **Figure 5H**, right). Both vectors showed comparable HA expression and spread within the dorsal striatum (**Figure 5I**). Female MCI-Park mice were excluded from surgical experiments due to their small body size (average body weight <12g) at P30.

Together, these findings show that AiE0450h-AADC-HA improves motor phenotypes in the progressive MCI-Park model in combination with L-DOPA, following both systemic and direct striatal delivery. In contrast, systemic CMV-AADC-HA delivery was associated with increased mortality under matched conditions, while direct striatal delivery produced behavioral improvements comparable to the enhancer-driven cell type-specific vector. These results also reveal sex-dependent differences in L-DOPA sensitivity required to achieve behavioral improvement.

### Retro-orbital administration of AADC in other relevant cell types differentially rescued specific motor behaviors in PD models

To determine whether AADC expression in other cell types implicated in PD could influence motor phenotypes, we generated enhancer-driven vectors targeting striatal cholinergic interneurons, dopaminergic neurons in the VTA/SNc, and brain-wide astrocytes. Previously validated enhancers AiE0743m_3xC2^48^, AiE0888m^65^, and AiE0387m_3xC1^50^ were used to target cholinergic neurons, dopaminergic neurons, and astrocytes respectively. Each construct incorporated the same HBB3-containing AADC-3xHA transgene cassette used in earlier vectors (**Figure 6A**).

**Figure 6:**
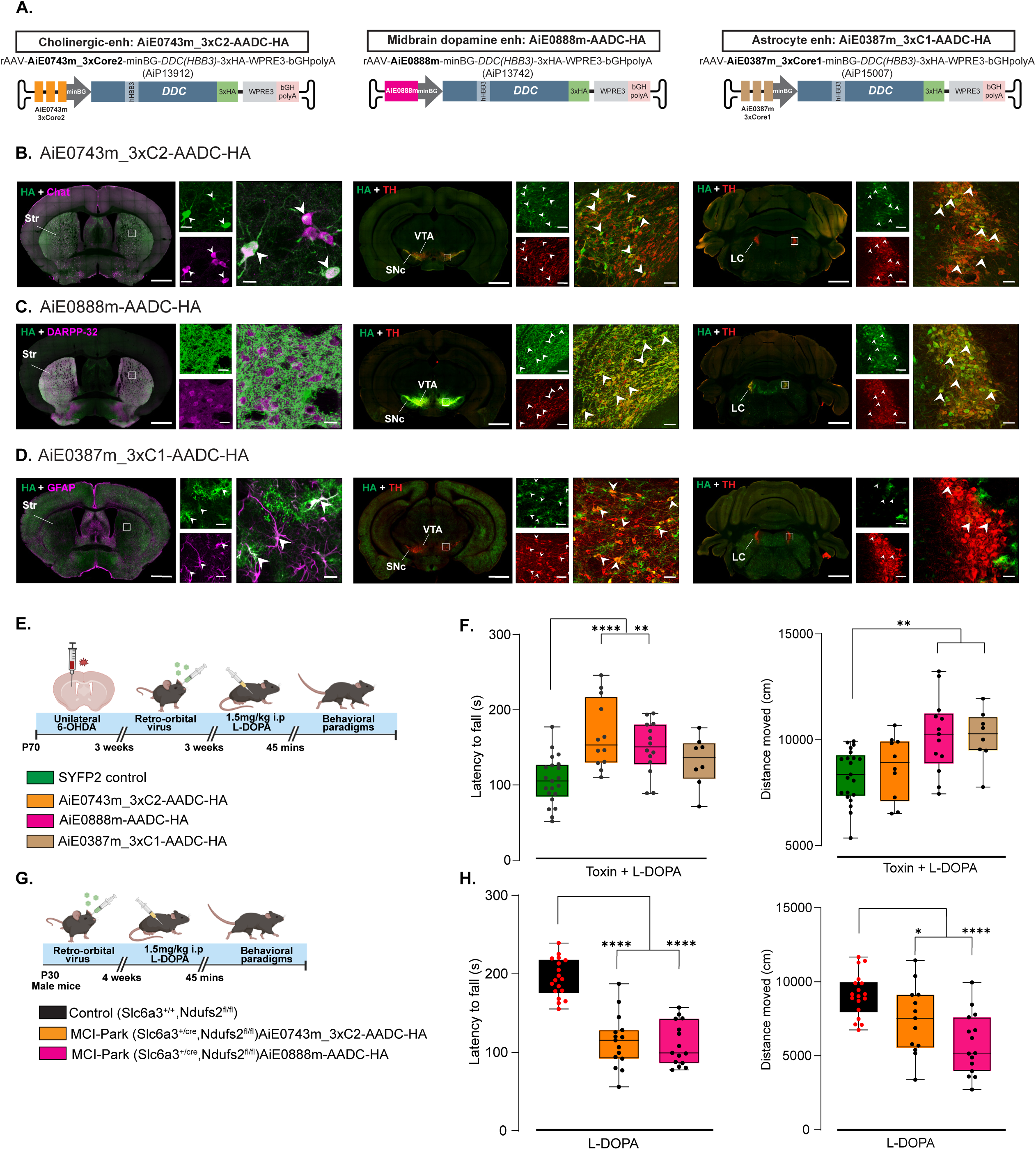
Retroorbital administration of AADC in other relevant cell types differentially rescued specific motor behaviors in PD model. (A) Schematic showing the vector designs for AiE0743m_3xC2-AADC-HA, AiE0888m-AADC-HA, and AiE0387m_3xC1-AADC-HA. (B) For AiE0743m_3xC2-AADC-HA: Representative images of striatal (AP = +0.14 to +0.26 from bregma) coronal images of HA (green) and Chat+ cells (magenta, left), midbrain (AP = -2.92 to -3.08 from bregma) coronal images of HA (green) and TH+ cells (red) in the middle panel, and LC (AP= -5.40 to -5.52 from bregma) coronal images of HA (green) and TH+ cells (red) in the right panel. Low magnification (10X) stitched, scale bars, 500 µm; High magnification (20X), scale bars, 10 µm for STR and 20 µm for VTA and LC. White arrows depict representative colocalized cells. (C) For AiE0888m-AADC-HA: Representative images of striatal (AP = +0.14 to +0.26 from bregma) coronal images of HA (green) and DARPP-32+ MSNs (magenta, left), midbrain (AP = -2.92 to -3.08 from bregma) coronal images of HA (green) and TH+ cells (red) in the middle panel, and LC (AP = -5.40 to -5.52 from bregma) coronal images of HA (green) and TH+ cells (red) in the right panel. Low magnification (10X) stitched, scale bars, 500 µm; High magnification (20X), scale bars, 10 µm for the striatum and 20 µm for VTA and LC. White arrows depict representative colocalized cells. (D) For AiE0387m_3xC1-AADC-HA: Representative images of striatal (AP = +0.14 to +0.26 from bregma) coronal images of HA (green) and GFAP+ cells (magenta, left), midbrain (AP = -2.92 to -3.08 from bregma) coronal images of HA (green) and TH+ cells (red) in the middle panel, and LC (AP = -5.40 to -5.52 from bregma) coronal images of HA (green) and TH+ cells (red) in the right panel. Low magnification (10X) stitched, scale bars, 500 µm; High magnification (20X), scale bars, 10 µm for the striatum and 20 µm for VTA and LC. White arrows depict representative colocalized cells. (E) Schematic overview of the injections and the behavioral tests. 6-OHDA toxin was stereotactically injected into the right dorsal striatum of P70 mice. 3 weeks after the surgery, they received RO injection of the viral vectors, and a series of behavioral tests were performed 3 weeks later, 45 mins after L-DOPA intraperitoneal injection. (F) Retention time of the mice in the accelerating rotarod test (left) and quantification of total distance traveled in the open field test (right). Data represented as box plots indicating 25^th^ to 75^th^ percentiles and median as the solid line with whiskers denoting minimum and maximum values. n = 21 (SYFP2 control), 8-14 (viral vectors). The *p* values were calculated by one-way ANOVA followed by Dunnett’s post hoc test. ∗∗*p* < 0.01, ∗∗∗∗*p* < 0.0001. (G) MCI-Park (*Slc6a3*^+/cre^; *Ndufs2*^fl/fl^) and age-matched littermate controls (*Slc6a3*^+/+^; *Ndufs2*^fl/fl^) were RO injected with the viral vectors at P30-35. 4 weeks later at around P60, a series of behavioral tests were performed 45 mins after L-DOPA intraperitoneal injection. (H) Retention time of the mice in the accelerating rotarod test (left) and quantification of total distance traveled in the open field test (right). Data represented as box plots indicating 25^th^ to 75^th^ percentiles and median as the solid line with whiskers denoting minimum and maximum values. n = 10 (WT), 13-15 (viral vectors). The *p* values were calculated by one-way ANOVA followed by Dunnett’s post hoc test. ∗*p* < 0.05, ∗∗∗∗*p* < 0.0001.

Vectors were packaged in PHP.eB and delivered RO (1e12 vg) into adult C57Bl6/J mice. Three weeks later, brain sections were immunostained with antibodies against HA and cell-type markers for cholinergic interneurons (choline acetyltransferase, ChAT^66^), dopaminergic neurons (TH), and astrocytes (glial fibrillary acidic protein, GFAP^67^) (**Figure 6B–D**).

The AiE0743m_3xC2 vector drove transgene-produced HA expression in striatal cholinergic neurons as expected (white arrows, **Figure 6B**, left) and also labeled TH+ neurons in the VTA, SNc, and LC (white arrows, **Figure 6B**, middle and right). Whereas the AiE0888m enhancer drove robust HA expression in dopaminergic fibers within the striatum (**Figure 6C**, left) and the target dopaminergic neurons of the VTA and SNc (white arrows, **Figure 6C**, middle). The vector also labeled TH+ neurons in the LC (white arrows, **Figure 6C**, right).

The AiE0387m_3xC1 enhancer drove robust HA expression in GFAP+ astrocytes throughout the brain, including the striatum, VTA, SNc, and LC (**Figure 6D**), consistent with the broader astrocyte expression previously observed with SYFP2 reporter (https://portal.brain-map.org/genetic-tools/genetic-tools-atlas).

To test whether AADC expression in these cell types could influence motor phenotypes, vectors were delivered RO (1e12 vg) to 6-OHDA–lesioned mice, followed by behavioral testing three weeks later after low-dose L-DOPA (1.5 mg/kg; **Figure 6E**).

Mice receiving AiE0743m_3xC2-AADC-HA (167.11 ± 13.34 s; p < 0.0001) or AiE0888m-AADC-HA (149.64 ± 9.23 s; p = 0.0029) showed significantly improved rotarod performance relative to SYFP2 controls (105.79 ± 6.97 s; **Figure 6F**, left). In contrast, AiE0387m_3xC1-AADC-HA did not significantly increase their latency to fall (131.54 ± 11.85 s; p = 0.2381). In the open-field assay, AiE0888m-AADC-HA (10210.70 ± 494.52 cm; p = 0.0014) and AiE0387m_3xC1-AADC-HA (10154.90 ± 449.68 cm; p = 0.0096) increased total distance traveled relative to controls (8291.98 ± 273.89 cm), whereas AiE0743m_3xC2-AADC-HA did not produce a significant change (8702.63 ± 471.64 cm; p = 0.8273; **Figure 6F**, right).

To determine whether these vectors could also influence motor phenotypes in the progressive MCI-Park model, AiE0743m_3xC2-AADC-HA and AiE0888m-AADC-HA were RO delivered (1e12 vg), and behavioral testing was performed four weeks later following 1.5 mg/kg L-DOPA (**Figure 6G**).

Unlike the unilateral toxin model, viral AADC expression in these non-MSN cell types did not improve motor phenotypes in MCI-Park male mice at the same low-dose L-DOPA. Compared with the controls (rotarod: 194.63 ± 5.73 s, open field: 9121.13 ± 346.46 cm), virally treated MCI-Park mice continued to show significantly impaired rotarod performance (AiE0743m_3xC2-AADC-HA: 114.02 ± 8.82 s; AiE0888m-AADC-HA: 110.64 ± 7.12 s; p < 0.0001) and reduced open-field activity (7420.88 ± 640.15 cm, p=0.0372 and 5791.73 ± 531.76 cm, p<0.0001 respectively, **Figure 6H**).

Together, these findings show that AADC expression in distinct neuronal and glial populations differentially influences motor phenotypes in the 6-OHDA toxin model. Midbrain dopaminergic targeting improved both rotarod and open-field performance, whereas striatal cholinergic interneuron targeting improved rotarod performance alone and astrocyte targeting selectively increased open-field activity. In contrast, only MSN-targeted AADC expression improved motor deficits in the progressive MCI-Park model, suggesting that striatal MSNs may preferentially support AADC-dependent dopamine production when substrate availability is limited. Collective behavioral and histological results are summarized in **Figure 6-supplement table 1 and 2** respectively.

## Discussion

Here we developed enhancer-driven AAV vectors expressing human AADC and characterized their expression and behavioral effects across multiple PD mouse models. Systemic delivery showed that AAV-AADC vectors maintained selective expression within the striatum while also labeling dopaminergic regions that endogenously express *Ddc*, such as the VTA/SNc and LC. In combination with low-dose L-DOPA, MSN-targeted AADC improved motor phenotypes in both toxin-induced and progressive genetic PD models and reduced dyskinesia-like behaviors in the 6-OHDA model following prolonged treatment. Comparison across select cell types further revealed that distinct cellular populations influence different behaviors in the toxin model, whereas only MSN-targeted expression improved motor deficits in the progressive MCI-Park model at the subthreshold L-DOPA dose.

Immunohistochemical analysis revealed that AADC labeling was observed not only within enhancer-defined striatal populations but also in TH⁺ neurons in dopaminergic regions. Extrastriatal expression was more pronounced following systemic delivery and markedly reduced after direct striatal injection, suggesting that viral entry mechanisms may determine the extent of spread beyond the target region, potentially via retrograde transport through axonal terminals^68,69^ rather than synaptic connectivity. Further studies are needed to determine the molecular basis of this extrastriatal expression at the protein, DNA, and RNA levels. However, while expression in residual dopaminergic neurons following systemic delivery likely contributed to behavioral improvement in PD models, it is notable that restriction of AADC expression predominantly to MSNs through intrastriatal administration remained sufficient to improve motor deficits under low-dose L-DOPA conditions.

PD patients require continued L-DOPA treatment, and chronic exposure frequently causes severe dyskinesia^70,71^, motivating efforts to mitigate the adverse effects of long-term therapy. The short plasma half-life of L-DOPA^72^, together with progressively longer off periods as disease advances^11,73^, produces pulsatile stimulation of dopamine receptors that alters basal ganglia output, resulting in LID^9,10,74,75^. In the 6-OHDA model, MSN-driven AADC reduced dyskinesia-like behaviors during prolonged L-DOPA exposure, suggesting that targeted AADC expression may stabilize dopamine production under fluctuating substrate availability. In addition, lower L-DOPA doses may achieve therapeutic benefit in the presence of AADC, potentially delaying dyskinesia onset. These findings are consistent with previous reports showing increased striatal dopamine levels following transduction of postsynaptic neurons in the putamen with ubiquitously expressed AADC^27,76^. Future measurements of dopamine levels following cell type-specific AADC delivery will further refine understanding of how distinct cellular populations contribute to behavioral rescue.

Although dopaminergic neurons constitute an intuitive target for AADC expression, their progressive degeneration in PD limits their therapeutic utility relative to AADC deficiency, where midbrain dopaminergic neurons and their projections remain structurally intact^23,77^. Despite this limitation, systemic enhancer-driven targeting of midbrain dopaminergic neurons improved motor performance in the toxin model, consistent with prior reports showing behavioral recovery following direct SN delivery of AADC to surviving dopaminergic neurons^64^. Emerging diagnostic approaches, including the FDA-approved α-synuclein seed assay^78^ and retinal thickness measurement^79,80^, may enable earlier detection of PD, potentially increasing the feasibility of dopaminergic neuron-targeted interventions. Nevertheless, the substantial loss of dopaminergic neurons in later PD stages underscores the need for alternative cell type-targeted strategies. In addition to MSNs, striatal cholinergic interneurons and astrocytes represent candidate targets due to their regulatory roles in striatal circuitry and L-DOPA metabolism^43,44,81–83^. In the 6-OHDA model, AADC expression in MSNs or cholinergic interneurons improved rotarod performance without affecting open-field activity, whereas astrocyte-targeted expression increased exploratory behavior without improving rotarod performance. This dissociation is consistent with the distinct behavioral domains captured by these assays^84–86^ and the established roles of these cell types, with cholinergic interneurons preferentially regulating motor coordination and balance measured on the accelerating rotarod^87^ and astrocytes modulating spontaneous locomotion in the open-field assay^88,89^. Similarly, distinct MSN subregions differentially regulate motor learning and locomotor activity, supporting their functional heterogeneity within striatal output pathways^90–92^. Together, these findings show that targeting different cell types can each support motor rescue in PD, with selective improvements aligned with their distinct circuit-specific functions.

In the MCI-Park model, however, only MSN-targeted AADC expression improved motor phenotypes under low L-DOPA conditions. Unlike the unilateral 6-OHDA model, this genetic model produces bilateral and progressive dopaminergic degeneration that more closely reflects the chronic loss of nigrostriatal input observed in PD. The selective efficacy of MSN targeting in this context suggests that striatal projection neurons may provide an attractive alternative for AADC-mediated therapy when dopaminergic neurons are substantially depleted. Furthermore, sex-dependent differences were observed in the MCI-Park model, where female mice required higher L-DOPA doses to reveal behavioral improvement following AADC expression. Differences in dopaminergic signaling and L-DOPA pharmacokinetics including various molecular and physiological variables like estrogen levels, L-DOPA clearance rate, urate levels^93–95^ have been reported previously and may contribute to this observation. These findings are consistent with clinical reports of faster disease progression, longer L-DOPA wearing-off periods, and greater dyskinesia risk in female patients^93,96^, highlighting the importance of developing tailored therapeutic strategies.

In summary, this work demonstrates the successful application of enhancer-driven, cell type-specific AADC expression to improve motor function in PD models while reducing L-DOPA–associated side effects. These vectors can be administered locally or paired with BBB-penetrant capsids for precise targeting, providing a framework for the development of next-generation cell-and circuit-informed gene therapies for PD.

## Data availability

All data, resources, and reagents will be made available upon request to the Lead Contact, Tanya L. Daigle. Mouse enhancer data are available at the Allen Genetic Tools Atlas (https://portal.brain-map.org/genetic-tools/genetic-tools-atlas). All AAV viral vector plasmids are freely available for research use at Addgene (addgene.org/)

## Key Resource Table

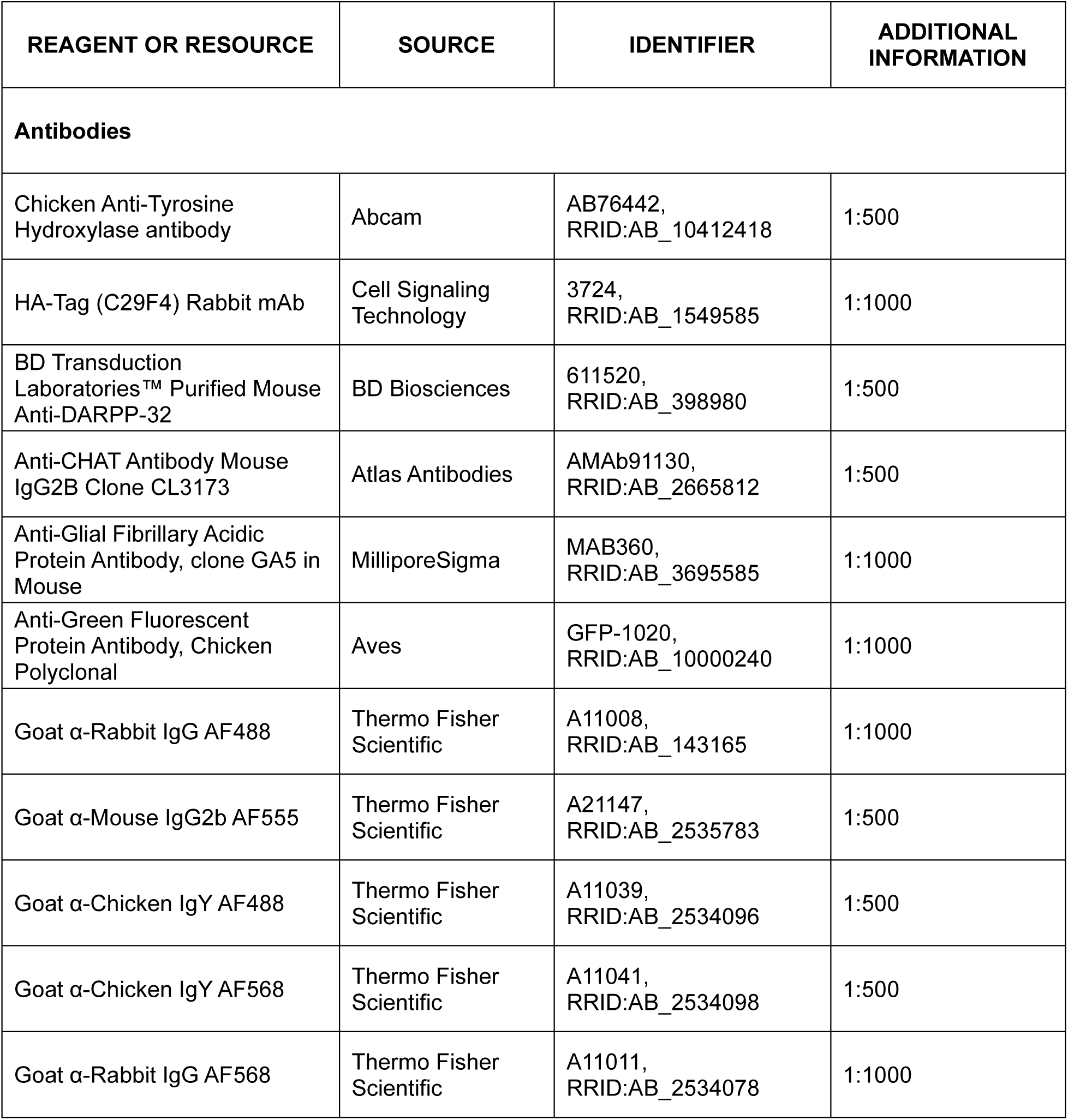
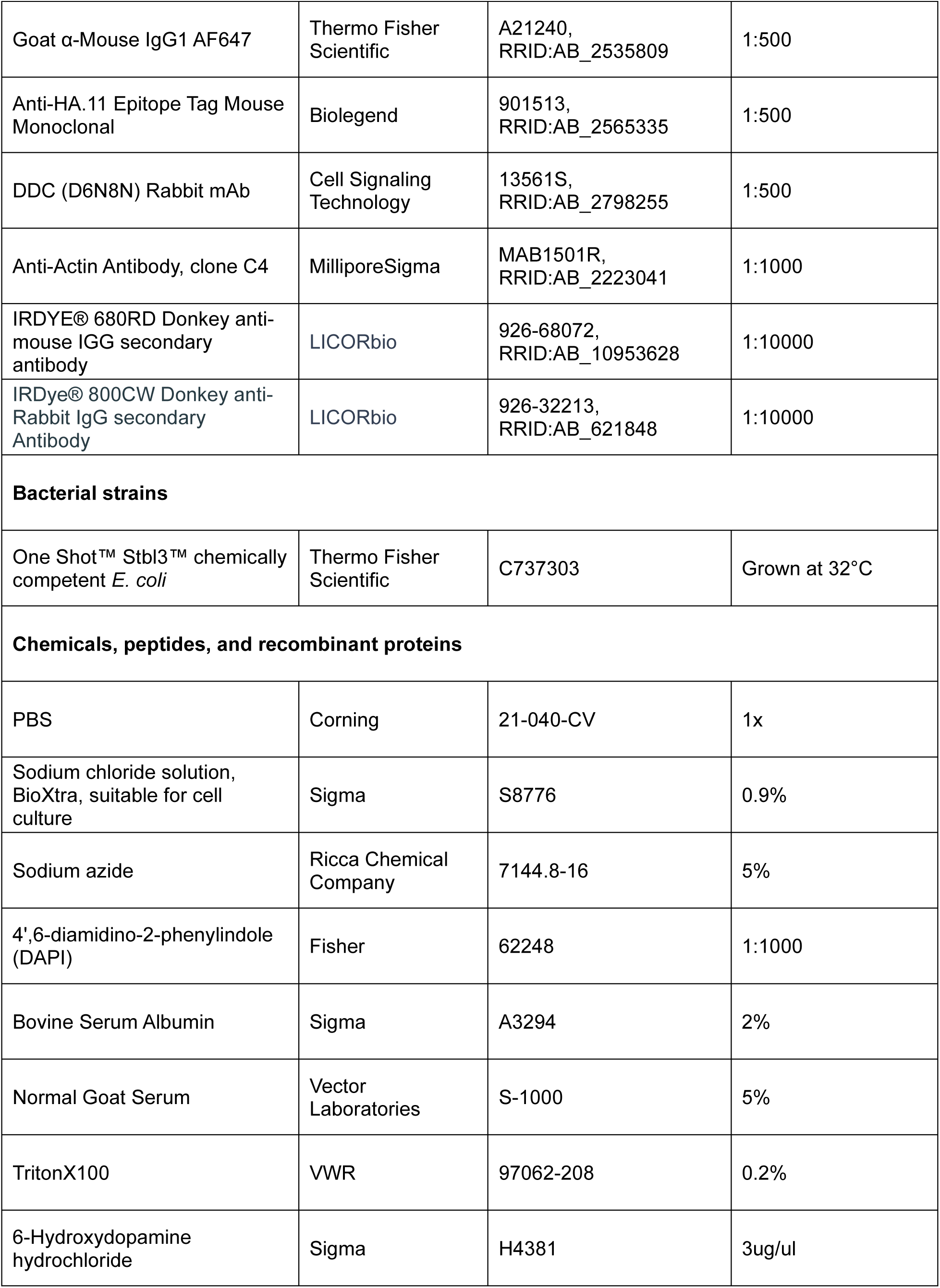
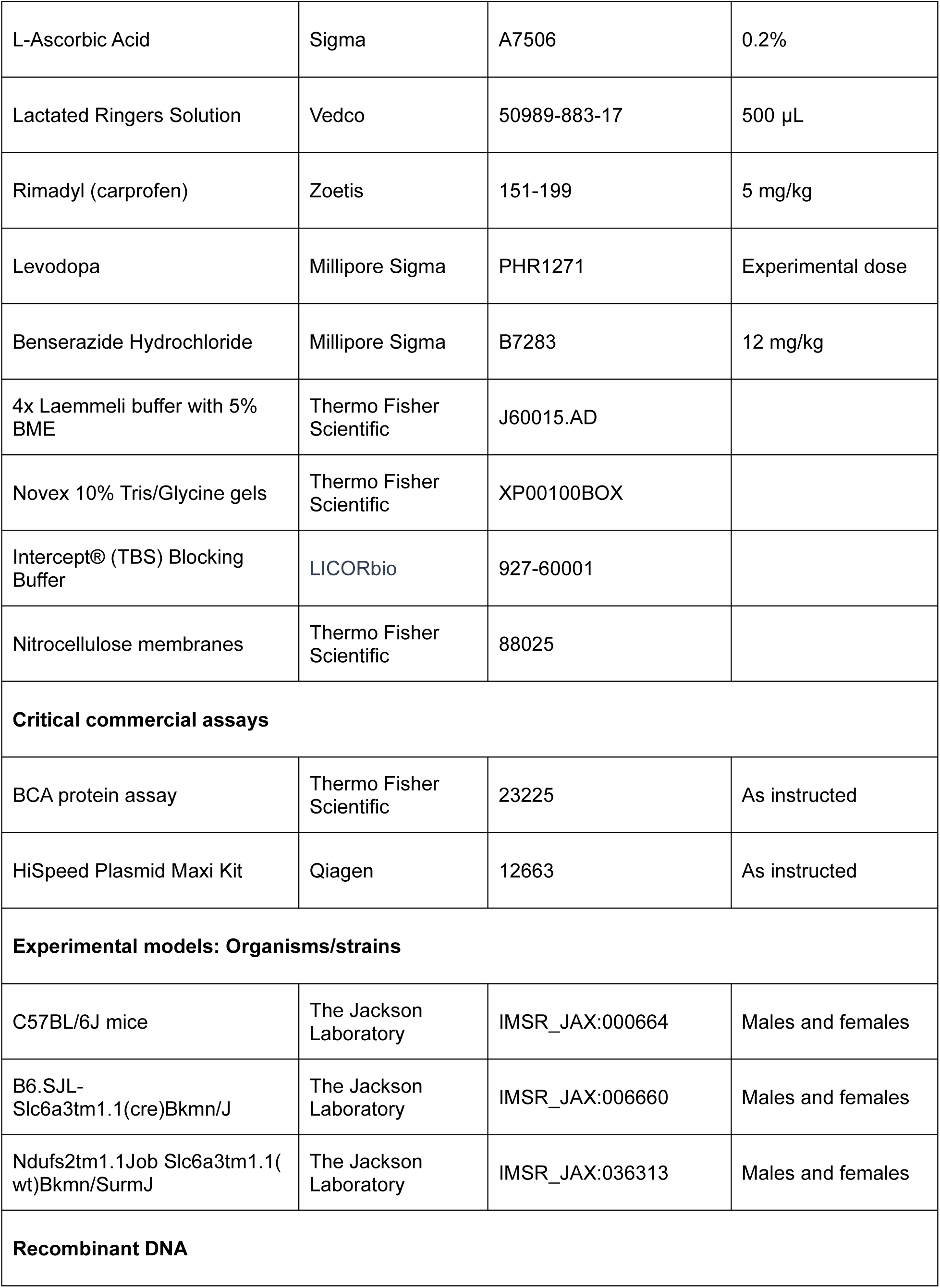
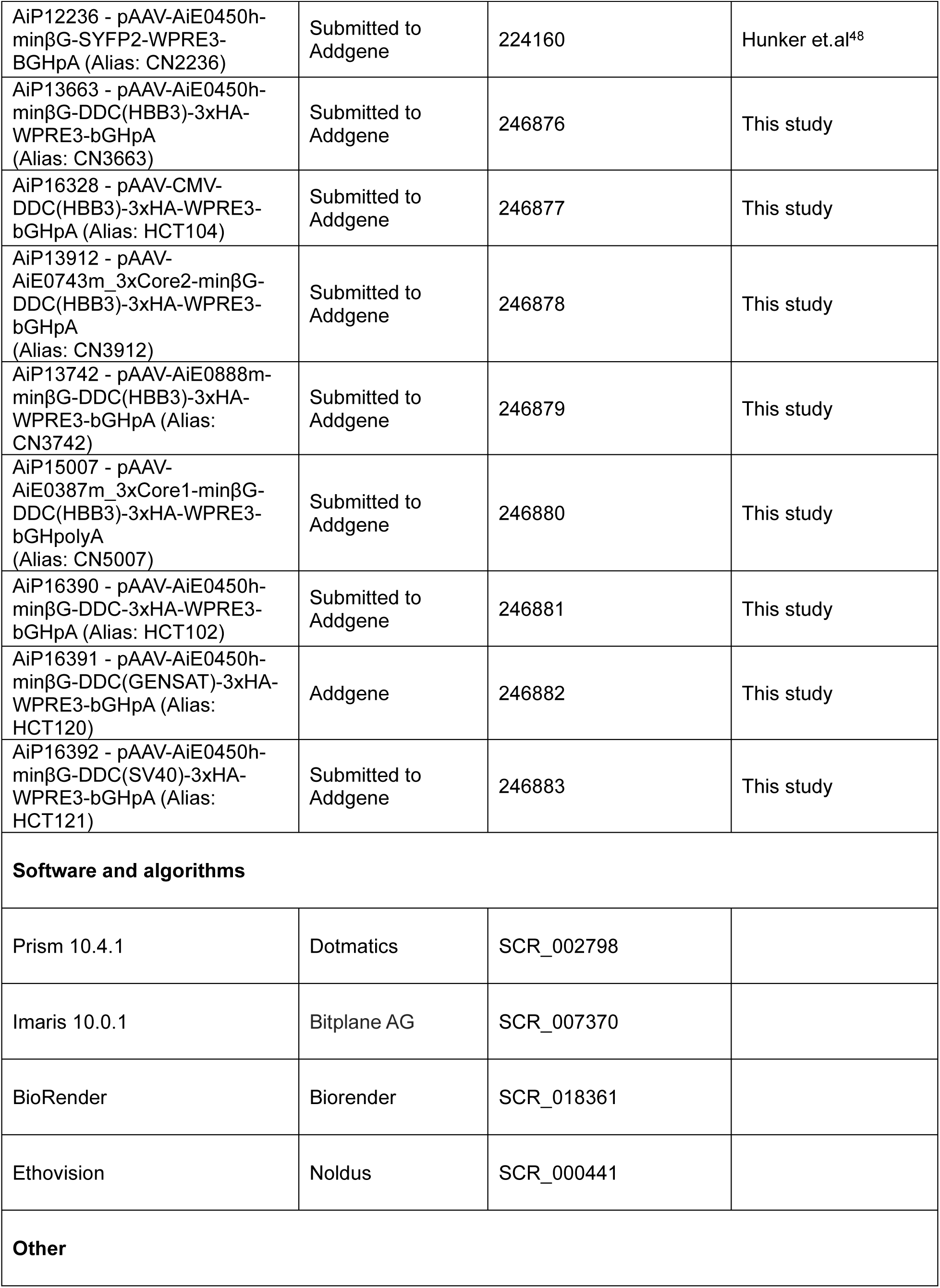
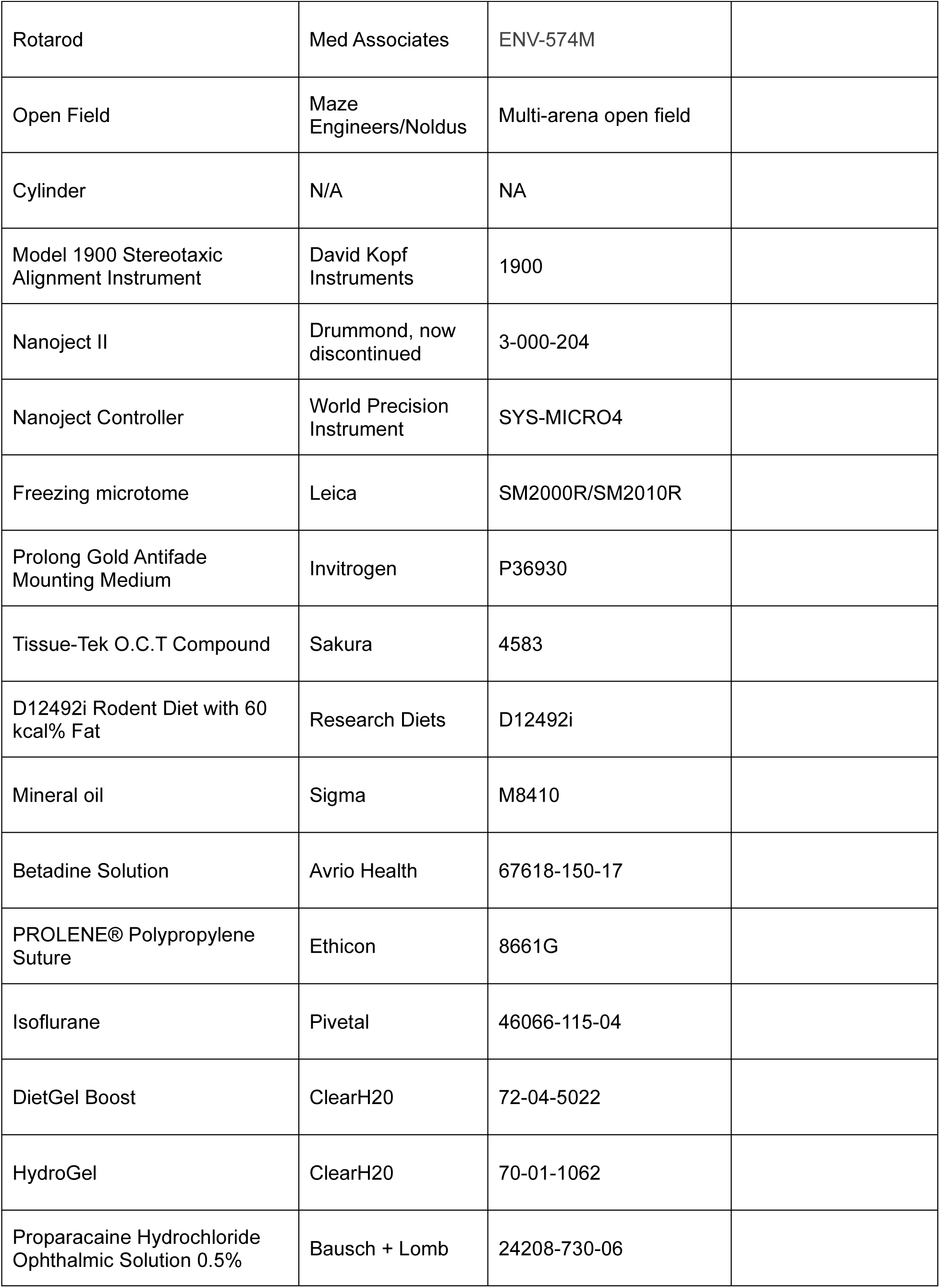

## Experimental model and subject details

### Mouse breeding and husbandry

For vector validations and the 6-OHDA toxin experiments, adult C57Bl/6J mice were bred and housed under Institutional Care and Use Committee (IACUC) protocols 2202 and 2505 at the Allen Institute, with no more than five animals per cage, maintained on a 12-hour light/dark cycle, with food and water provided *ad libitum*. Enrichment, such as plastic shelter and nesting materials, were provided. Adult C57BL/6J mice were also purchased directly from The Jackson Laboratory (Strain #000664, RRID: IMSR_JAX:000664) as needed and housed at the Allen Institute under the same conditions. Both male and female mice were used for experiments, and the minimal number of animals was used for each experimental group. Sample sizes were established following the prevailing standards in behavioral neuroscience research for mice and our previous experience. All experiments and analysis described in this study were conducted and randomized double-blindly. Animals with anophthalmia or microphthalmia or other obvious conditions (e.g. penile prolapse) were excluded from experiments.

Mice carrying homozygous floxed *Ndufs2* (*Ndufs2*^fl/fl^) were crossed with double heterozygous mice expressing Cre recombinase (Cre) under the control of the DA transporter (DAT, encoded by *Slc6a3*) promoter (*Slc6a3*^+/cre^) as well as one floxed allele of *Ndufs2 (Ndufs*2^+/fl^) on a C57BL/6J background. This breeding scheme allowed us to generate both MCI-Park mice (*Slc6a3*^+/cre^; *Ndufs*2^fl/fl^) and age-matched littermate controls expressing floxed *Ndufs2* in the absence of Cre recombinase (*Slc6a3*^+/+^; *Ndufs2*^fl/fl^) while deriving future double heterozygous breeders. To create our in-house parent lines, heterozygous *Slc6a3-cre* mice (B6.SJL-*Slc6a3^tm1.1(cre)Bkmn^*/J; Strain#:006660, RRID:IMSR_JAX:006660) and homozygous floxed *Ndufs2 mice (Ndufs2^tm1.1Job^ Slc6a3^tm^1.1(wt)Bkmn/SurmJ;* Strain#:036313, RRID:IMSR_JAX:036313) were obtained from the Jackson Laboratories, bred and housed under IACUC protocols 2202 and 2505 at the Allen Institute.

## Methods details

### Viral vector construction

All recombinant AAV vectors (rAAV) backbones contained inverted terminal repeats (ITRs) required for viral packaging, a woodchuck hepatitis virus posttranscriptional regulatory element (WPRE3) to enhance expression, and the bGH polyadenylation signal for transcript stability. The previously reported rAAV-AiE0450h-minBG-SYFP-WPRE2-bGHpolyA (AiP12236) served as the initial template^48^.

For vectors encoding *DDC* (AiP13663, AiP16328, AiP13912, AiP13742, AiP15007, AiP16392, AiP16391, AiP16390), the open reading frame was optimized using the Integrated DNA Technologies (IDT) Codon Optimization Tool followed by additional rounds of manual editing to facilitate gBlock synthesis and improve stability. These refinements included reducing GC-rich domains, eliminating long or repetitive elements, and removing predicted cryptic splice donor/acceptor motifs, TATA-like sequences, homopolymeric WWWWWW tracts, and CpG-rich regions.

For constructs that contained an intron (AiP13663, AiP16328, AiP13742, AiP15007, AiP16392, AiP16391), the intron was inserted at the first high-confidence splice donor/acceptor site (CAG^G) within the open reading frame to promote higher expression. All plasmids were generated using standard molecular cloning techniques and were amplified in One Shot™ Stbl3™ chemically competent *E. coli* (Thermo Fisher Scientific, Cat. #C737303) at 32°C, purified using the HiSpeed Plasmid Maxi Kit (Qiagen, Cat. #12663) and validated by full plasmid sequencing with Plasmidsaurus® (Oxford Nanopore platform).

### Viral packaging and titering

Iodixanol-gradient purified rAAVs of the PHP.eB serotypes (AiE0450h-AADC(no intron)-HA; AiP16390, AiE0450h-AADC(SV40 small-t)-HA; AiP16391, AiE0450h-AADC(SV40)-HA; AiP16392) were generated in house using previously described methods^45,47^. The average titer of these viral preps was 5e13 vg/mL (ranging from 3.92-7.78e13 vg/mL) and were quantified using droplet digital PCR (QX 200, Bio-Rad) by targeting the ITR sequences. The following primers and probes were used: forward primer 5′-GGAACCCCTAGTGATGGAGTT-3′, reverse primer 5′-CGGCCTCAGTGAGCGA-3′, and probe 5′-/56-FAM/CGCGCAGAG/ZEN/AGGGAGTGG/3IABkFQ/-3′.

AiE0450h-SYFP2; AiP12236, AiE0450h-AADC-HA; AiP13663, CMV-AADC-HA; AiP16328, AiE0743m_3xC2-AADC-HA; AiP13912, AiE0888m-AADC-HA; AiP13742, and AiE0387m_3xC1-AADC-HA; AiP15007 purified viruses were packaged by PackGene Biotech at the titer of 2e13 vg/mL, quantified using qPCR with the same primer set as above.

### Mouse retro-orbital (RO) injections of AAVs

Mice were briefly anesthetized with isoflurane and 1e12 vg were delivered into the right venous sinus in a volume of 90µL via RO injection. 0.5% proparacaine hydrochloride ophthalmic solution (Bausch+Lomb, Cat# 24208-730-06) was then applied to the injected eye. Tissue samples were collected for analysis 3-4 weeks post-injection.

### Mouse stereotactic surgeries

#### 6-OHDA toxin surgeries

Pre-operative care started a week prior to surgery, where supplementary food was provided in a limited amount to both vehicle-control and toxin-lesion mice (P70) to increase their body weights to ≥ 28 g on surgery date. Supplementary foods consisted of DietGel Boost (Clear H2O, 370 Kcal/g) and D12492i Rodent Diet (Research Diets, 60% kcal fat). This supplementary food was discontinued upon recovery by 10-14 days (weight stabilization post-surgery).

Mice were anesthetized with 4% isoflurane and maintained at 1.5-2% in a stereotaxic frame (Kopf 1900) equipped with a heating pad (37 °C) to maintain their body temperature. At the start of the surgery, all mice were injected subcutaneously with 500 μL of LRS (Lactated Ringers Solution, Vedco) and 5 mg/kg carprofen (Rimadyl, Zoetis, Cat# 151-199). Ophthalmic ointment (Systane, Alcon) was applied to the eyes to prevent corneal drying. The lesion was induced in right dorsal striatum by injecting 1.5 μL of 6-OHDA hydrochloride (H4381, Sigma Aldrich) at a concentration of 3 µg/µL dissolved in sterile 1x PBS with 0.2% ascorbic acid (A7506, Sigma Aldrich), according to the following coordinates (mm from bregma): anteroposterior (AP) +0.26; mediolateral (ML) +2.00; and dorsoventral (DV) −3.4 (from skull). Control mice received a sham lesion, consisting of unilateral injections of 1.5 μL of vehicle (0.2% ascorbic acid in sterile 1x PBS). The rate of infusion was maintained at 150 nL/min with a Nanoject II Nanoliter Injector (Drummond, Cat#3-000-204) and Micro4 Controller (World Precision Instruments, SYS-MICRO4), and after infusion, the glass capillary was kept at the injection site for 5 minutes before being withdrawn slowly to avoid efflux. Post injection, the mice were returned to the home cage kept on a heating pad (37°C) to recover for an hour before returning them to vivarium.

#### Intra-striatal viral surgeries

The injections were performed bilaterally in the dorsal striatum, according to the following coordinates (mm from bregma): anteroposterior (AP) +0.26; mediolateral (ML) +/- 2.00 and dorsoventral (DV) −3.4 (from skull). Viral vectors were thawed and diluted 1:10 with sterile 1x PBS solution to reach 2e12 vg. 500 nL (final 1e9 vg) was delivered to both the hemispheres at the rate of 100 nL/min while undiluted vector (2e13 vg) was injected for the high dose (final 1e10 vg) experiment. The remainder of the procedure was performed the same as before.

### Mouse behavioral phenotyping

#### Acclimation

Adult mice (P60-100, depending on the PD model) were used for behavioral testing. All tests were conducted in the light phase. Mice were placed in the testing room with reduced light density to acclimate for at least 30 minutes in home cages before each paradigm. All tests were run on different days and counterbalanced to avoid the inter-behavioral effect as well as possible L-DOPA interaction. The testing equipment was cleaned with Peroxigard (Virox Technologies Inc.) and/or 70% ethanol and air dried between mice.

#### L-DOPA preparation

Levodopa (Sigma Aldrich, PHR1271) was used to prepare the L-DOPA solution (in 0.9% sterile saline) at the doses mentioned in the paper (1.5 mg/kg to 20 mg/kg, depending on the experiment). A peripheral decarboxylase inhibitor, Benserazide (12 mg/kg, B7283, Sigma Aldrich), was always added to the preparation to inhibit the enzyme DOPA decarboxylase in the body’s periphery, increasing the L-DOPA content in the brain^97^. L-DOPA/Benserazide drug solution should be made fresh daily before injections, if possible, or stored in light-protected condition at 4 °C up to 48 hours. The mice were always placed in behavioral apparatus 45 mins post-L-DOPA/Benserazide intraperitoneal administration, either after a single injection or on the last day of the prolonged application.

#### Accelerating Rotarod

Rotarod accelerating from 4–40 rotations per min over 300 s was used (ENV-574M, Med Associates) to test their motor behavior. Mice were run for three trials with 3 min inter-trial interval. Each trial ended if mice fell off the rod, started rotating with the rod for one whole rotation, or successfully stayed on rod for the full 300 s. The rods, walls, and fall containers were cleaned with 70% ethanol and air dried before and after each trial. The behavioral performance (latency to fall) was quantified as the average of the three trials.

#### Open Field

General locomotor behavior of freely moving mice was evaluated in multi-unit open field boxes (EthoVision XT, Noldus), each box measuring 40 x 40 x 30 cm. The mice were video recorded for 30 minutes during which the body center point of each mouse was live tracked and total distance traveled was compared using EthoVision XT (Noldus, RRID:SCR_000441) software.

#### Cylinder test

A clear acrylic (McMaster-Carr) cylinder of 20 cm (diameter) and 30 cm (height) was used to evaluate the frequency of paw touches during explorative activity, to observe and quantify asymmetric behaviors like rotations, dragging of feet, and falling on one side, and to score abnormal involuntary movements in LID. The test apparatus was placed in front of a mirror to observe the mice from all angles. Every session was video recorded for 10 minutes to be analyzed by an experimenter blind to the groups, and the values were randomly checked by another blinded experimenter. The room was dimly lit during the entire trial, and the walls and floor of the cylinder were cleaned with Peroxigard (Virox Technologies Inc.) and air dried between mice.

#### Contralateral paw touches

Forelimb use during supported rearing was assessed by scoring the number of load-bearing contacts on the cylinder wall. Individual contact was counted when the mouse returned to a resting position and reared again to touch the cylinder wall. Only the initial contact was marked as ipsilateral first or contralateral first or both simultaneously, regardless of whether they used or took off the second paw afterwards. Ipsilateral and contralateral were defined relative to the 6-OHDA-lesioned side. Percent contralateral paw touches were calculated as [contralateral first) /total observations] * 100.

#### Contralateral rotations

Videos recorded in the cylinder were further analyzed to quantify the total number of ipsilateral and contralateral turns to the lesion over 10 minutes. 180-degree turns were scored as one, while 360-degree turns were scored as two^98,99^. Net contralateral turns were calculated as [total contralateral - total ipsilateral].

#### Abnormal involuntary movements (AIMs)

Videos recorded in the cylinder were used to assess abnormal involuntary movements (AIMs) as defined in previous studies^7,98,100^. The AIMs were classified into three different categories based on their topographic distribution: (i) axial AIMs-twisting of the neck and upper body towards the side contralateral to the lesion. Unlike the rotations described earlier, these were characterized by the absence of full body turns. (ii) orolingual AIMs-empty jaw movements, lip smacking, and tongue protrusion; (iii) forelimb AIMs-brisk, purposeless movements of the contralateral forelimb, sometimes combined with empty grabbing movement or tapping of the paw. Instead of separating the categories into subtypes based on AIM severity scale^100^, the occurrences were evaluated as presence and absence of the dyskinetic phenotype. Each individual incidence was counted from start of an event to end of the event when the mouse returned to a resting position and classified into the three categories defined earlier. Though rare, if within one incidence, more than one category of AIMs occurred, each would be counted separately. The hyperkinetic and dystonic movements had to be involuntary, insistent, and clearly distinguished from naturally occurring behaviors like grooming, sniffing, rearing and gnawing for them to be considered in quantification. Total occurrence of AIMs was calculated by adding the number of incidences in the three defined types over 10 mins.

#### Immunostaining

Mice were transcardially perfused with 4% PFA (paraformaldehyde) in 1x PBS. After perfusion, tissues were kept in the fixation solution overnight at 4°C, then transferred to 30% sucrose solution for 24 hours. Using a freezing microtome (Leica SM2000R, RRID:SCR_018456), tissue was mounted on OCT medium (Tissue-Tek® O.C.T. Compound Cat# 4583), sectioned (50 μm thickness) and collected free-floating.

Sections were washed three times in 1x PBS at room temperature and were blocked in 1x PBS containing 5% normal goat serum (Vector Laboratories, Cat# S-1000-20), 0.2% Triton X-100 (Millipore Sigma, Cat# X100), 2% bovine serum albumin (Millipore Sigma, Cat# A9418-5G) for 1 hour at room temperature. Sections were then incubated overnight at 4°C in the same blocking solution containing primary antibodies: Chicken Anti-Tyrosine Hydroxylase antibody (1:500; AB76442, Abcam, RRID:AB_10412418), HA-Tag (C29F4) Rabbit mAb (1:1000; 3724, Cell Signaling Technology, RRID:AB_1549585), Purified Mouse Anti-DARPP-32 (1:500; 611520, BD Biosciences, RRID:AB_398980), Anti-CHAT Antibody Mouse IgG2B Clone CL3173 (1:500; AMAb91130, Atlas Antibodies, RRID:AB_2665812), Anti-Glial Fibrillary Acidic Protein Antibody, clone GA5 in Mouse (1:1000; MAB360, Millipore Sigma, RRID:AB_3695585). After primary incubation, sections were washed three times in 1x PBS and incubated in the same blocking solution containing DAPI (1:1000; Fisher Scientific, Cat# 62248) and secondary antibodies: Goat α-Rabbit IgG AF488 (1:1000; A11008, Thermo Fisher Scientific, RRID:AB_143165), Goat α-Mouse IgG12B AF555 (1:500; A21147, Thermo Fisher Scientific, RRID:AB_2535783), Goat α-Chicken IgY AF568 (1:500; A11041, Thermo Fisher Scientific, RRID:AB_2534098), Goat α-Rabbit IgG AF568 (1:1000; A11011, Thermo Fisher Scientific, RRID:AB_2534078), Goat α-Mouse IgG1 AF647 (1:500; A21240, Thermo Fisher Scientific, RRID:AB_2535809) for 2 hours at room temperature. Following secondary incubation, sections were washed three times in 1x PBS. All steps were performed under gentle agitation. Stained tissue was mounted onto Superfrost slides (Fisher Scientific, Cat# 12-550-15) with ProLong Gold antifade reagent (Invitrogen, Cat# P36930, RRID:SCR_015961).

#### Imaging

Images were acquired with a Nikon Eclipse Ti2 epifluorescence microscope and a Zeiss LSM 880 confocal microscope using manufacturer’s software. Whole section images were acquired with a 10x objective using the same intensity and exposure time across samples at a resolution of 512×512. Z-stacks were collected in 6μm increments and collapsed into maximum intensity projections using manufacturer’s software. Higher magnification images were acquired with a 20x objective using the same intensity and exposure time across samples at a resolution of 1024×1024. Z-stacks were collected in 1 μm increments and collapsed into maximum intensity projections using Imaris software (Bitplane 10.0.1, RRID:SCR_007370). Brightness, contrast, and pseudo-coloring were matched across images using Imaris tools. For quantification, the brightness and contrast were kept uniform across all images and spots were identified for individual channels using the automated Imaris spot detection (10 μm) with background subtraction applied. Colocalization was assigned automatically with a constant shortest distance to spots (<10 μm). Percent completeness of AADC-HA labeling was calculated by taking the number of double-positive cells (i.e., HA+ and TH+) and dividing that by the total number of TH+ cells. Percent specificity was calculated by taking the number of double-positive cells (i.e., HA+ and TH+) and dividing that number by the total number of virally labeled cells (i.e., HA+).

#### Western Blot

Mice were euthanized without anesthetics by cervical dislocation followed by rapid decapitation. Dissected brain regions were flash-frozen in liquid nitrogen and stored at -80°C until processing. Samples were sonicated in 200 uL of 95 °C 1% SDS with protease inhibitors (Thermo Fisher Scientific, Cat#78441), using previously described methods^101^. Protein levels were assessed using a BCA kit (Thermo Fisher Scientific, Cat#23225) and lysate was denatured by combining with 4x Laemmeli buffer containing 5% BME and then boiled at 95°C for 5 mins. 50 ug of protein was loaded into 10% Tris/Glycine gels (Thermo Fisher Scientific, Cat#XP00100BOX) and then transferred onto nitrocellulose membranes (Thermo Fisher Scientific, Cat#88025) for 2.5 hours at 25V and at 4°C. Membranes were blocked in TBS with 5% BSA for 1 hour at room temperature, then incubated overnight at 4°C with primary antibodies in TBS containing 0.05% Tween-20 and 5% BSA. Primary antibodies included anti-HA (1:500; Biolegend, Cat#901513, RRID:AB_2565335), anti-AADC (1:500; Cell Signaling Cat#13561S, RRID:AB_2798255), and anti-actin (1:1000; Millipore, Cat#MAB1501R, RRID: AB_2223041. After washing in TBS with 0.1% Tween-20, membranes were incubated with fluorescent secondary antibodies (1:10,000 each; Ms 680 and Rb 800; LI-COR, Cat#926-68072, RRID:AB_10953628 and 926-32213, RRID:AB_621848) in LI-COR TBS blocking buffer (Cat#927-60001), washed, and imaged using an Odyssey Infrared Imaging System (LI-COR). Protein signals were quantified using LI-COR analysis software and normalized to actin.

### Statistical Analyses

Statistical analyses were conducted using GraphPad Prism 10.4.1 (Dotmatics/GraphPad Software, RRID:SCR_002798) and graphical illustrations were created using BioRender (RRID:SCR_018361). When the data is represented as box plots, the boxes indicate 25th to 75th percentiles and median as the solid line with whiskers denoting minimum and maximum values. When the data is represented as bar graphs, the error bars indicate the standard error of the mean (SEM). Sample size (n) refers to biological replicates unless otherwise noted in the figure legends. All behavioral datasets were independently replicated at least twice and plotted together. Histological results were independently confirmed. Statistical significance was assessed by Welch’s t-test, Mann-Whitney, and one-way ANOVA, where appropriate, followed by the post-hoc tests as indicated in the figure legends. Normality was tested by Shapiro-Wilk and Kolmogorov-Smirnov tests. The level of significance was set at p<0.05. Data points deviating more than 2 SD values from the mean were excluded from the analyses.

## Acknowledgements

We thank members of the Allen Institute animal care team for providing animal care, surgical support, and assistance with tissue collection for the study. We thank Dr. Rui Costa for kindly sharing the MCI-Park transgenic lines and Dr. D. James Surmeier for critical comments on the manuscript. Many schematics were created with BioRender.com. Research reported in the publication was solely supported by the Paul G. Allen Foundation. We thank the Allen Institute founder, Paul G. Allen, for his vision, encouragement, and support.

## Author contributions

T.L.D, J.T.T. and A.C. designed the study. J.K.M., X.O-A., R.M., and A.C.H. designed and constructed vectors. A.C., A.F., N.T., V.O., and M.N.L. performed histology and analyses, M.A.Q. performed biochemistry and analyses. A.F., A.C., M.D., and E.K., performed injections, behavior, and analyses. M.R., E.L., L.S., D.R., C.H., B.C., M.L., O.H., T.H., T.W., N.U., C.B., and J.B. performed perfusions, dissections, and provided mouse health support. D.N., R.K., S.K., N.D., and S.Y. performed in-house viral production. K.R., and V.W. performed genetic model breeding, colony maintenance, genotyping, and provided mouse health support. K.G. provided program management support. T.L.D, J.T.T., E.S.L., B.P.L., and G.D.H. provided leadership. A.C. and T.L.D. wrote the manuscript with support from many co-authors.

---

**Figure 1–supplement figure 1:**
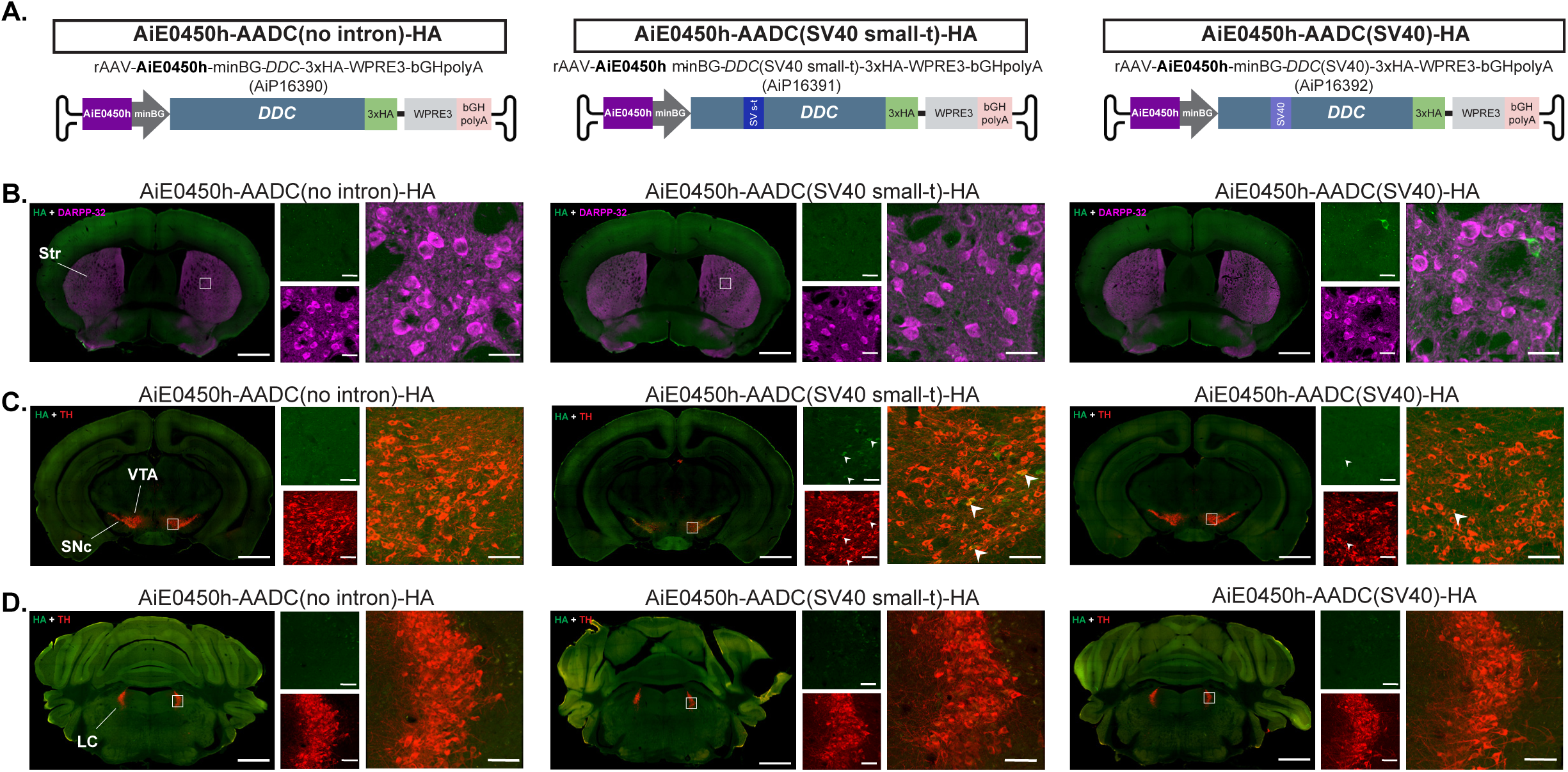
Development and characterization of different AADC transgene designs. (A) Schematic showing the vector designs for AiE0450h-AADC-HA with no intron, SV40 small-t intron, or longer SV40 intron. (B) Representative striatal (AP = +0.14 to +0.26 from bregma) coronal images of HA (green) and DARPP-32+ MSNs (magenta). (C) Representative midbrain (AP = -2.92 to -3.08 from bregma) and (D) representative LC (AP= -5.40 to -5.52 from bregma) coronal images of HA (green) and TH (red), for AiE0450h-AADC(no intron)-HA (left), AiE0450h-AADC(SV40 small-t)-HA (middle), and AiE0450h-AADC(SV40)-HA (right), following 1e12 vg RO administration. Low magnification (10X) stitched, scale bars, 500 μm; High magnification (20X), scale bars, 10 μm for the striatum and 20 μm for VTA and LC. White arrows depict representative colocalized cells.

**Figure 1–supplement figure 2:**
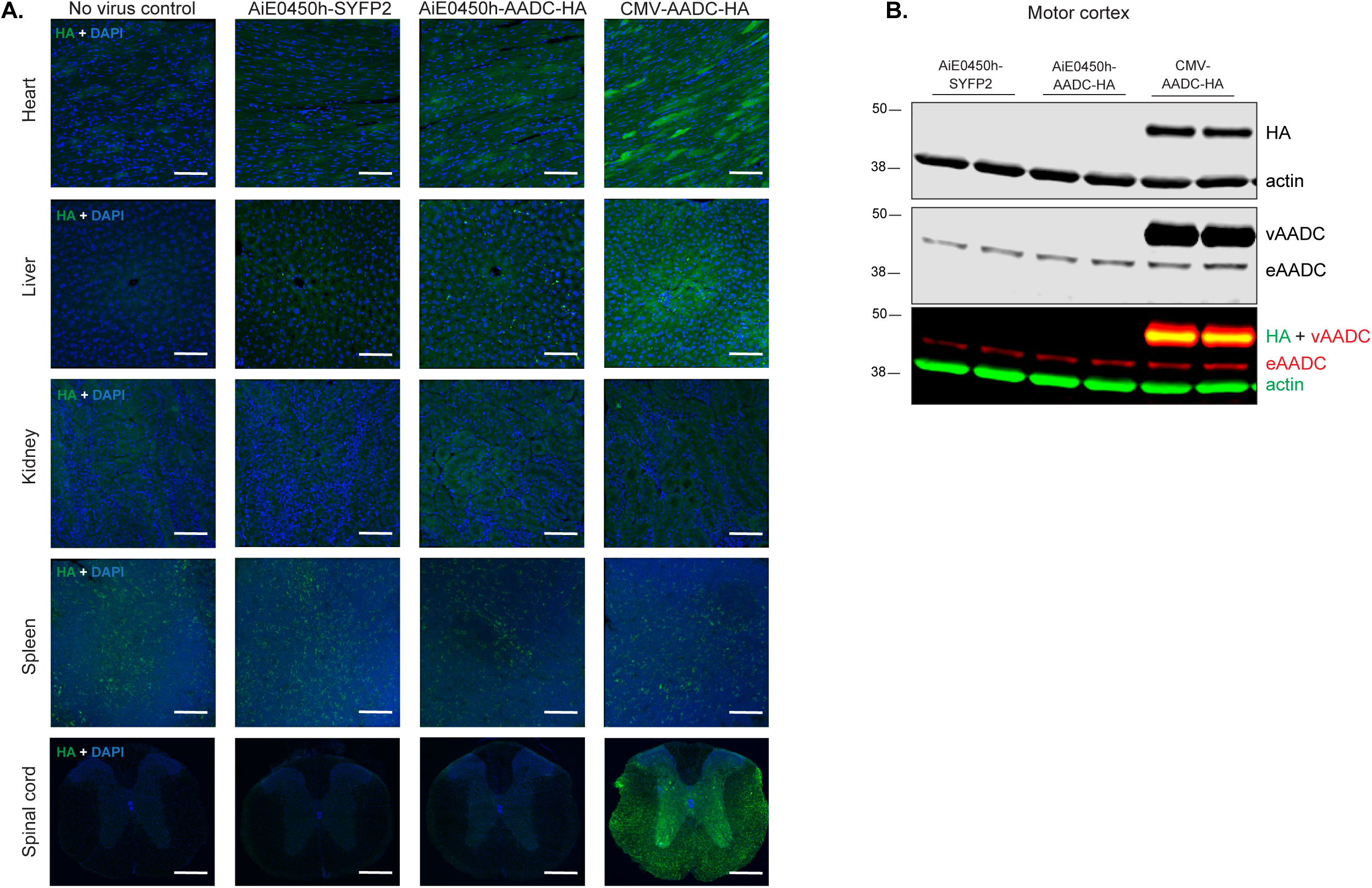
Biodistribution of viral transgene in CNS and peripheral organs. (A) Peripheral organ and spinal cord biodistribution data for no-virus control, AiE0450h-SYFP2, AiE0450h-AADC-HA, and CMV-AADC-HA following 1e12 vg RO administration. Representative confocal images of HA (green) or DAPI (blue) in transverse sections prepared from heart, liver, kidney, spleen, and spinal cord of control and virally treated mice. High magnification (20X), scale bars, 20 μm. (B) Representative western blot of motor cortex-enriched tissue punches from virally treated mice showing HA and actin bands (top), AADC bands (middle), and merged image (bottom) with HA (green) and viral AADC (vAADC; red) overlaying. The endogenous AADC (eAADC) appears as a lower-molecular-weight band. Actin (green) was used as a loading control.

**Figure 2-supplement figure 1:**
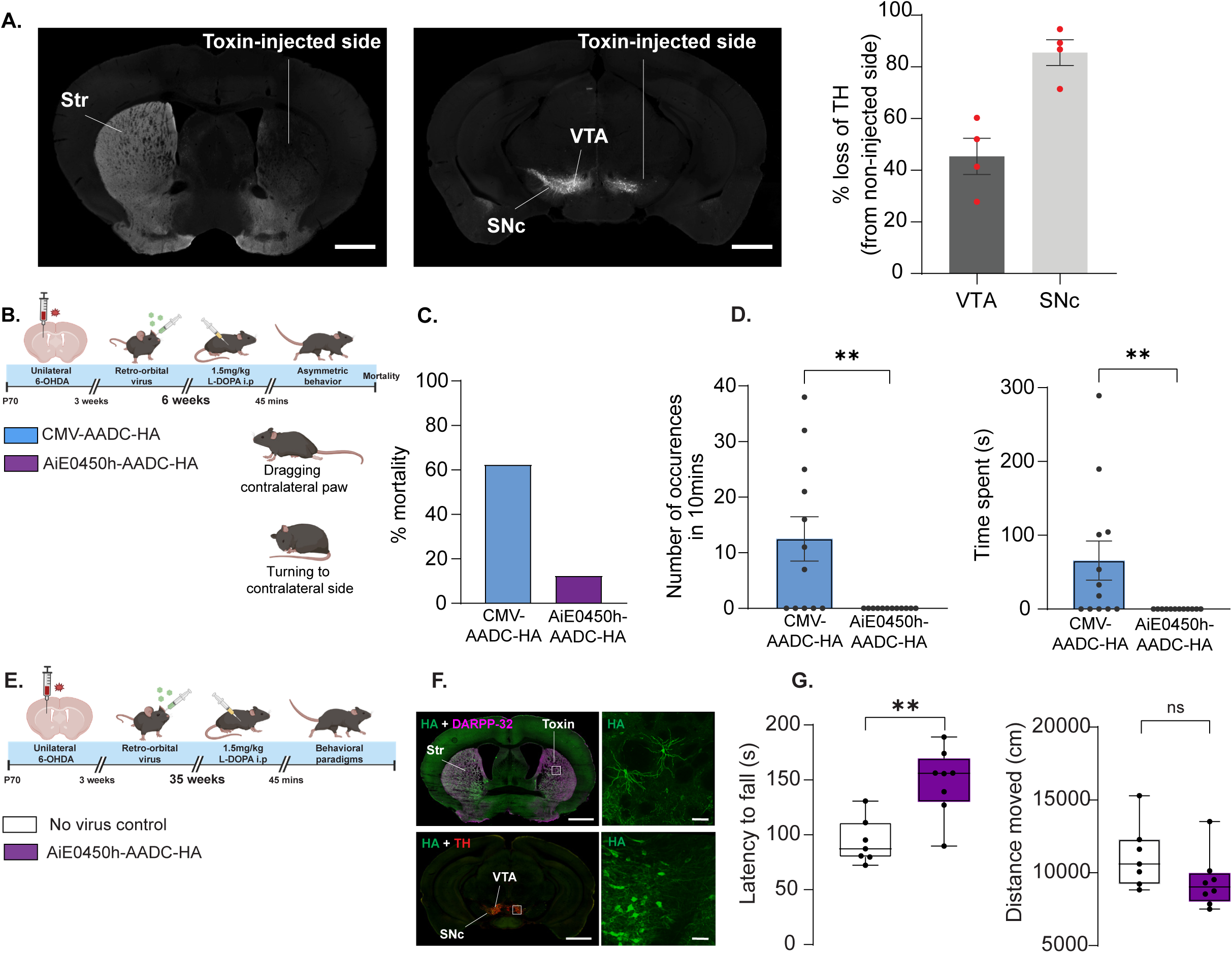
6-OHDA model validation and phenotypic effect of AADC-AAVs (at 6 weeks and 35 weeks) (A) Representative coronal images of the striatum and midbrain demonstrating the unilateral depletion of TH staining (gray) after toxin administration. Low magnification (10X) stitched, scale bars, 500 µm. Quantification of the loss of TH+ cells in VTA and SNc (% reduction compared to the non-injected hemisphere). Data represented as mean ± SEM. n = 4 mice, each data point average of 3 sections. (B) Schematic overview of the injections and the behavioral tests to access side effects at 6 weeks post-RO in toxin-lesioned mice. (C) Mortality rate 6 weeks after RO injection of the viral vectors (% out of total injected). (D) Quantification of contralateral asymmetric movements in a cylinder over 10 mins. Data represented as mean ± SEM. n = 12 per group. The p values were calculated by Mann-Whitney test, normality tested with Shapiro-Wilk test. **p < 0.01. (E) Schematic overview of the injections and the behavioral tests to access durability of enhancer-AADC at 35 weeks post-RO in toxin-lesioned mice. (F) Representative striatal and midbrain coronal images of HA (green) and DARPP-32+ MSNs (magenta) or TH+ cells (red) for AiE0450h-AADC-HA 35 weeks after RO injection in toxin-lesioned mice. Low magnification (10X) stitched, scale bars, 500 μm; High magnification (20X), scale bars, 10 μm for the striatum and 20 μm for VTA. (G) Retention time of the toxin-lesioned mice in the accelerating rotarod test (left) and total distance traveled in the open field test (right) 35 weeks after RO. Data represented as box plots indicating 25th to 75th percentiles and median as the solid line with whiskers denoting minimum and maximum values. n = 7-8 per group. The p values were calculated by Mann-Whitney test, normality tested with Shapiro-Wilk test. *p < 0.05, **p < 0.01, ***p < 0.001.

**Figure 3-supplement figure 1:**
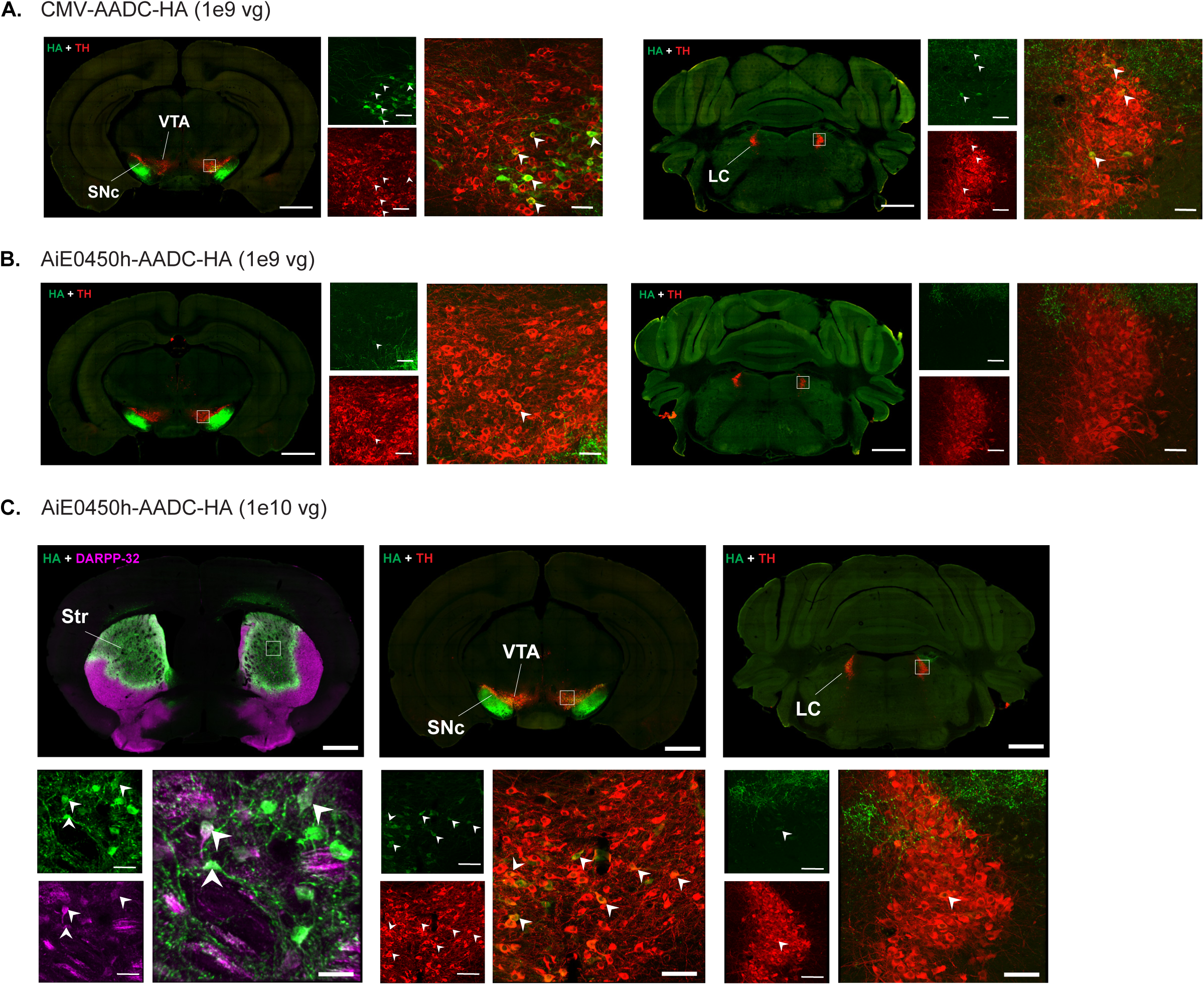
Extrastriatal expression of viral HA upon intrastriatal delivery of AADC-AAVs at two different titers. Representative midbrain (AP = -2.92 to -3.08 from bregma) and LC (AP = -5.40 to -5.52 from bregma) coronal images of HA (green) and TH+ cells (red) for (A) CMV-AADC-HA and (B) AiE0450h-AADC-HA at intrastriatal administration of the vector at behaviorally relevant 1e9 vg titer. Low magnification (10X) stitched, scale bars, 500 µm; High magnification (20X), scale bars, 20 µm. White arrows depict representative colocalized cells. (C) Representative coronal images of striatum (AP = +0.14 from bregma) for AiE0450h-AADC-HA showing HA (green) and DARPP-32+ MSNs (magenta), midbrain (AP = -3.08 from bregma), and LC (AP = -5.40 from bregma) showing HA (green) and TH+ cells (red), following at higher titer of 1e10 vg. Low magnification (10X) stitched, scale bars, 500 μm, top; High magnification (20X), scale bars, 10 μm for striatum (bottom left) and 20 μm for VTA (bottom middle) and LC (bottom right). White arrows depict representative colocalized cells.

**Figure 5-supplement figure 1:**
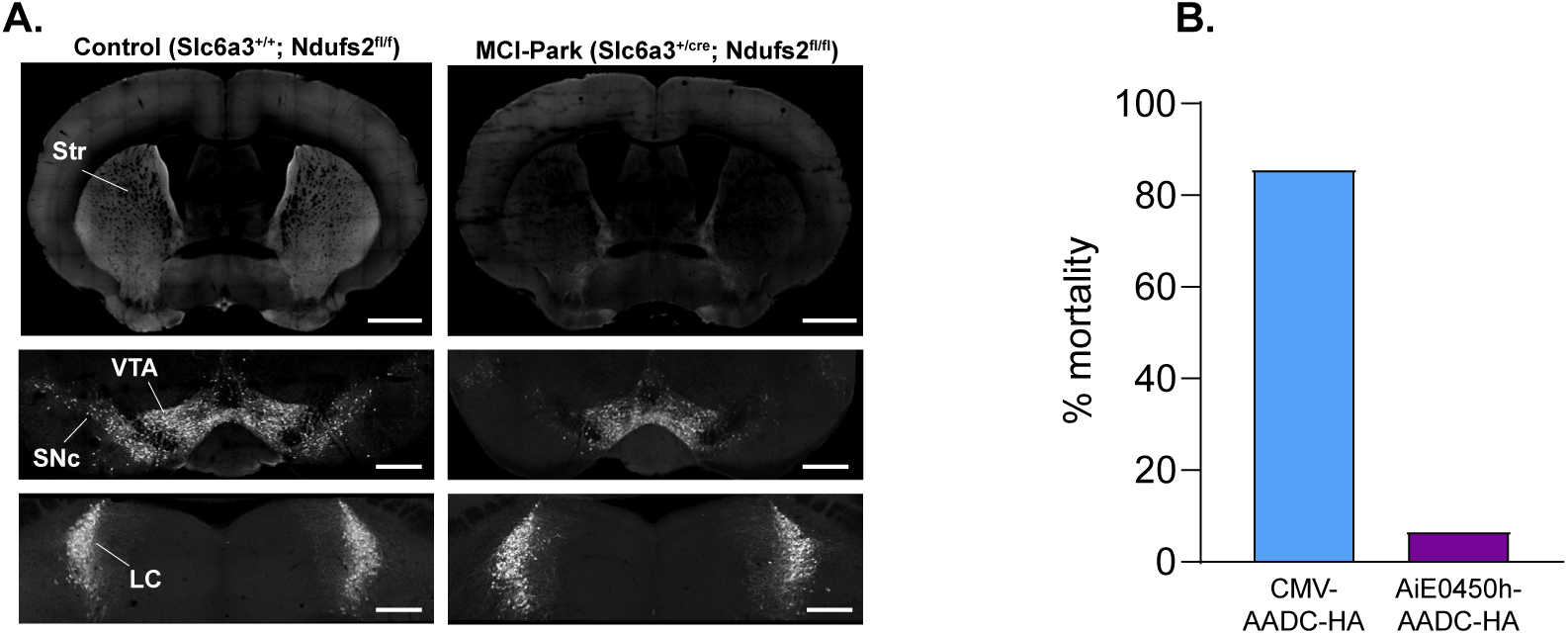
Validation of MCI-Park genetic model and phenotypic observation. (A) Representative coronal images of the striatum (top) and midbrain (bottom) demonstrating the bilateral depletion of TH staining (gray) in MCI-Park (Slc6a3+/cre; Ndufs2fl/fl, right) compared to the age-matched littermate controls (Slc6a3+/+; Ndufs2fl/fl, left). Low magnification (10X) stitched, scale bars, 500 μm. (B) Mortality rate 4 weeks after retroorbital injection of the viral vectors (% out of total injected).

**Figure 6-supplement table1:**
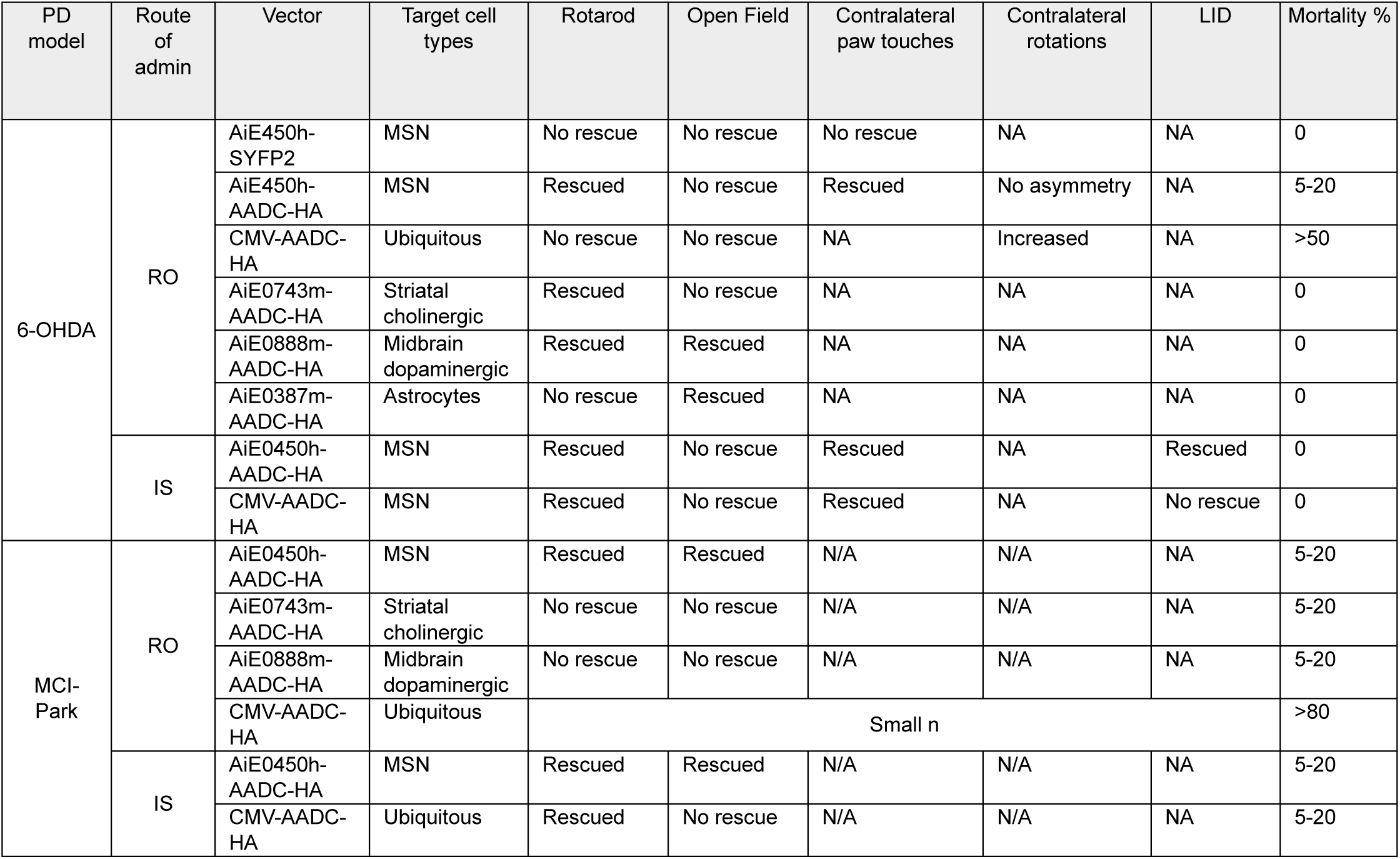
Summary table of behavioral results and mortality rates observed in 6-OHDA and MCI-Park models with AAVs delivered via two routes of administration. Rescued at 1.5 mg/ kg L-DOPA= statistically significant reversal of behavioral deficit with AAV treatment. No rescue = no change in the behavioral deficit with AAV treatment. No asymmetry = AAV did not produce this behavioral effect. Increased = AAV produced notable rotations. NA = data is not available and N/A = not applicable for the model

**Figure 6-supplement table2:**
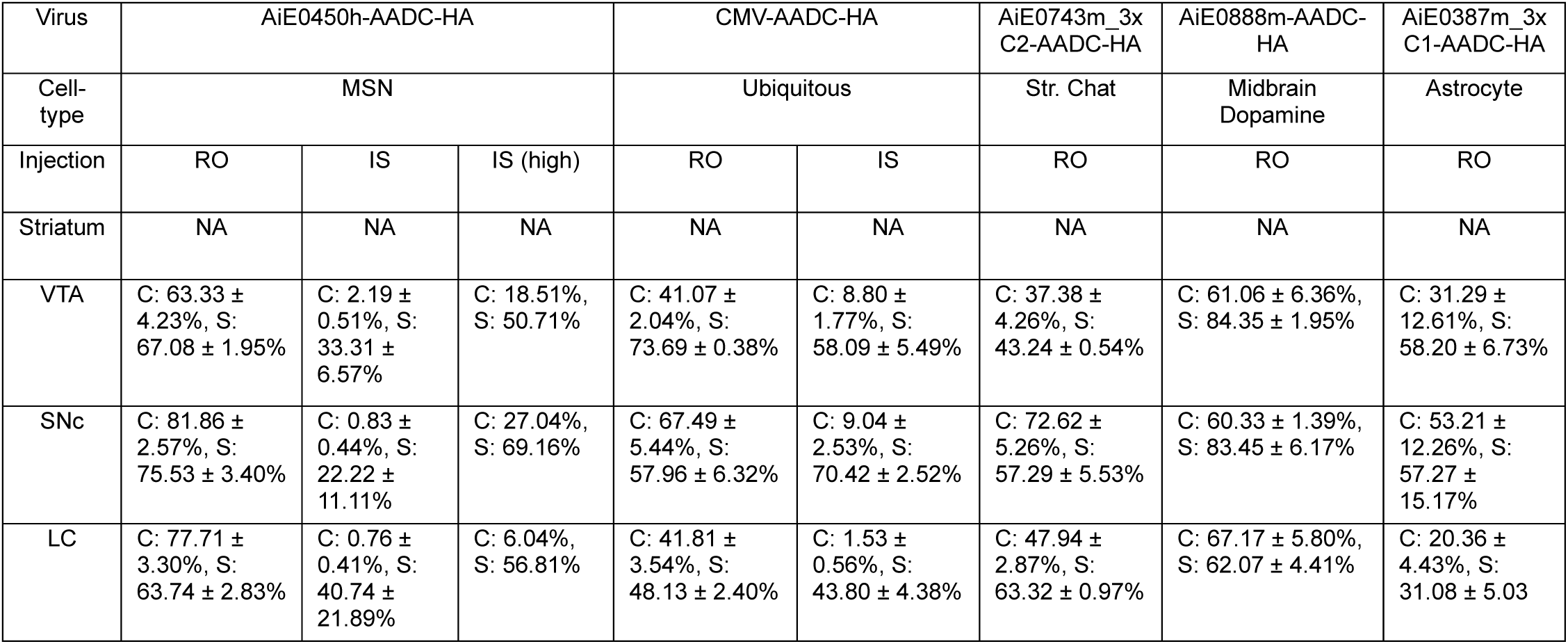
Summary table of histology data with AAVs delivered via two routes of administration. C: % Completeness of HA labeling in TH+ neurons (HA+ and TH+/ TH+). S: % Specificity of TH+ labeling within HA+ neurons (HA+ and TH+/HA+). NA = not analyzed due to dense labeling of fibers. Percentage values are the mean ± SEM from 3 sections of n = 3 mice each in all analyses except AiE0450h-AADC-HA IS (high) where it is mean of 3 sections from n= 1 mouse.

## References

1. DeLong, M. R. & Wichmann, T. Circuits and Circuit Disorders of the Basal Ganglia. Arch Neurol 64, 20 (2007).

2. Luk, K. C. et al. Pathological α-Synuclein Transmission Initiates Parkinson-like Neurodegeneration in Nontransgenic Mice. Science 338, 949–953 (2012).

3. Zhou, Z. D., Yi, L. X., Wang, D. Q., Lim, T. M. & Tan, E. K. Role of dopamine in the pathophysiology of Parkinson’s disease. Transl Neurodegener 12, 44 (2023).

4. Jankovic, J. Parkinson’s disease: clinical features and diagnosis. J Neurol Neurosurg Psychiatry 79, 368 (2008).

5. Moustafa, A. A. et al. Motor symptoms in Parkinson’s disease: A unified framework. Neuroscience & Biobehavioral Reviews 68, 727–740 (2016).

6. Bezard, E., Brotchie, J. M. & Gross, C. E. Pathophysiology of levodopa-induced dyskinesia: Potential for new therapies. Nat Rev Neurosci 2, 577–588 (2001).

7. Lundblad, M., Picconi, B., Lindgren, H. & Cenci, M. A. A model of l-DOPA-induced dyskinesia in 6-hydroxydopamine lesioned mice: relation to motor and cellular parameters of nigrostriatal function. Neurobiology of Disease 16, 110–123 (2004).

8. Lundblad, M. et al. Pharmacological validation of a mouse model of l-DOPA-induced dyskinesia. Experimental Neurology 194, 66–75 (2005).

9. Dwi Wahyu, I., Chiken, S., Hasegawa, T., Sano, H. & Nambu, A. Abnormal Cortico-Basal Ganglia Neurotransmission in a Mouse Model of l-DOPA-Induced Dyskinesia. J. Neurosci. 41, 2668–2683 (2021).

10. Yang, K. et al. Circuit Mechanisms of L-DOPA-Induced Dyskinesia (LID). Front. Neurosci. 15, 614412 (2021).

11. Kwon, D. K., Kwatra, M., Wang, J. & Ko, H. S. Levodopa-Induced Dyskinesia in Parkinson’s Disease: Pathogenesis and Emerging Treatment Strategies. Cells 11, 3736 (2022).

12. Byrne, B. J. et al. Current clinical applications of AAV-mediated gene therapy. Molecular Therapy 33, 2479–2516 (2025).

13. Merola, A. et al. Gene Therapy in Movement Disorders: A Systematic Review of Ongoing and Completed Clinical Trials. Front. Neurol. 12, 648532 (2021).

14. Naso, M. F., Tomkowicz, B., Perry, W. L. & Strohl, W. R. Adeno-Associated Virus (AAV) as a Vector for Gene Therapy. BioDrugs 31, 317–334 (2017).

15. Ogbonmide, T. et al. Gene Therapy for Spinal Muscular Atrophy (SMA): A Review of Current Challenges and Safety Considerations for Onasemnogene Abeparvovec (Zolgensma). Cureus https://doi.org/10.7759/cureus.36197 (2023) doi:10.7759/cureus.36197.

16. Reape, K. Z. & High, K. A. Trial by “Firsts”: Clinical Trial Design and Regulatory Considerations in the Development and Approval of the First AAV Gene Therapy Product in the United States. Cold Spring Harb Perspect Med 13, a041312 (2023).

17. Zhu, M. Y., Juorio, A. V., Paterson, I. A. & Boulton, A. A. Regulation of Aromatic L -Amino Acid Decarboxylase by Dopamine Receptors in the Rat Brain. Journal of Neurochemistry 58, 636–641 (1992).

18. Ciesielska, A. et al. Depletion of AADC activity in caudate nucleus and putamen of Parkinson’s disease patients; implications for ongoing AAV2-AADC gene therapy trial. PLoS ONE 12, e0169965 (2017).

19. Muramatsu, S. et al. A Phase I Study of Aromatic L-Amino Acid Decarboxylase Gene Therapy for Parkinson’s Disease. Molecular Therapy 18, 1731–1735 (2010).

20. Chien, Y.-H. et al. Efficacy and safety of AAV2 gene therapy in children with aromatic L-amino acid decarboxylase deficiency: an open-label, phase 1/2 trial. The Lancet Child & Adolescent Health 1, 265–273 (2017).

21. Kojima, K. et al. Gene therapy improves motor and mental function of aromatic l-amino acid decarboxylase deficiency. Brain 142, 322–333 (2019).

22. Hwu, W.-L. et al. Gene Therapy for Aromatic L -Amino Acid Decarboxylase Deficiency. Sci. Transl. Med. 4, (2012).

23. Pearson, T. S. et al. Gene therapy for aromatic L-amino acid decarboxylase deficiency by MR-guided direct delivery of AAV2-AADC to midbrain dopaminergic neurons. Nat Commun 12, 4251 (2021).

24. UpstazaTM Granted Marketing Authorization by European Commission as First Disease-Modifying Treatment for AADC Deficiency.

25. Golikeri, A., Yi, S. & Fashoyin-Aje, L. Eladocagene Exuparvovec for Aromatic L-Amino Acid Decarboxylase Deficiency. JAMA 333, 1449 (2025).

26. Tai, C.-H. et al. Long-term efficacy and safety of eladocagene exuparvovec in patients with AADC deficiency. Molecular Therapy 30, 509–518 (2022).

27. Sánchez-Pernaute, R., Harvey-White, J., Cunningham, J. & Bankiewicz, K. S. Functional Effect of Adeno-associated Virus Mediated Gene Transfer of Aromatic L-Amino Acid Decarboxylase into the Striatum of 6-OHDA-Lesioned Rats. Molecular Therapy 4, 324–330 (2001).

28. Sanftner, L. M. et al. Striatal Delivery of rAAV-hAADC to Rats with Preexisting Immunity to AAV. Molecular Therapy 9, 403–409 (2004).

29. Cunningham, J. et al. Biodistribution of Adeno-associated Virus Type-2 in Nonhuman Primates after Convection-enhanced Delivery to Brain. Molecular Therapy 16, 1267–1275 (2008).

30. Hadaczek, P. et al. Eight Years of Clinical Improvement in MPTP-Lesioned Primates After Gene Therapy With AAV2-hAADC. Molecular Therapy 18, 1458–1461 (2010).

31. Nutt, J. G. et al. Aromatic L-Amino Acid Decarboxylase Gene Therapy Enhances Levodopa Response in Parkinson’s Disease. Movement Disorders 35, 851–858 (2020).

32. Christine, C. W. et al. Safety and tolerability of putaminal *AADC* gene therapy for Parkinson disease. Neurology 73, 1662–1669 (2009).

33. Mittermeyer, G. et al. Long-Term Evaluation of a Phase 1 Study of AADC Gene Therapy for Parkinson’s Disease. Human Gene Therapy 23, 377–381 (2012).

34. Christine, C. W. et al. Magnetic resonance imaging–guided phase 1 trial of putaminal *AADC* gene therapy for Parkinson’s disease. Annals of Neurology 85, 704–714 (2019).

35. Christine, C. W. et al. Safety of AADC Gene Therapy for Moderately Advanced Parkinson Disease: Three-Year Outcomes From the PD-1101 Trial. Neurology 98, (2022).

36. Qin, J. Y. et al. Systematic Comparison of Constitutive Promoters and the Doxycycline-Inducible Promoter. PLoS ONE 5, e10611 (2010).

37. Gray, S. J. et al. Optimizing Promoters for Recombinant Adeno-Associated Virus-Mediated Gene Expression in the Peripheral and Central Nervous System Using Self-Complementary Vectors. Human Gene Therapy 22, 1143–1153 (2011).

38. Bäck, S. et al. Neuronal Activation Stimulates Cytomegalovirus Promoter-Driven Transgene Expression. Molecular Therapy - Methods & Clinical Development 14, 180–188 (2019).

39. Hwu, P. W. et al. Gene therapy in the putamen for curing AADC deficiency and Parkinson’s disease. EMBO Mol Med 13, e14712 (2021).

40. Ugrumov, M. V. Non-dopaminergic neurons partly expressing dopaminergic phenotype: Distribution in the brain, development and functional significance. Journal of Chemical Neuroanatomy 38, 241–256 (2009).

41. Kozina, E. A., Kim, A. R., Kurina, A. Y. & Ugrumov, M. V. Cooperative synthesis of dopamine by non-dopaminergic neurons as a compensatory mechanism in the striatum of mice with MPTP-induced Parkinsonism. Neurobiology of Disease 98, 108–121 (2017).

42. Troshev, D. et al. Striatal Neurons Partially Expressing a Dopaminergic Phenotype: Functional Significance and Regulation. IJMS 23, 11054 (2022).

43. Inyushin, M. Y. et al. L-DOPA Uptake in Astrocytic Endfeet Enwrapping Blood Vessels in Rat Brain. Parkinson’s Disease 2012, 1–8 (2012).

44. Asanuma, M., Miyazaki, I., Murakami, S., Diaz-Corrales, F. J. & Ogawa, N. Striatal Astrocytes Act as a Reservoir for L-DOPA. PLoS ONE 9, e106362 (2014).

45. Graybuck, L. T. et al. Enhancer viruses for combinatorial cell-subclass-specific labeling. Neuron 109, 1449–1464.e13 (2021).

46. Mich, J. K. et al. Functional enhancer elements drive subclass-selective expression from mouse to primate neocortex. Cell Reports 34, 108754 (2021).

47. Ben-Simon, Y. et al. A suite of enhancer AAVs and transgenic mouse lines for genetic access to cortical cell types. Cell 188, 3045–3064.e23 (2025).

48. Hunker, A. C. et al. Enhancer AAV toolbox for accessing and perturbing striatal cell types and circuits. Neuron 113, 1507–1524.e17 (2025).

49. Kussick, E. et al. Enhancer AAVs for targeting spinal motor neurons and descending motor pathways in rodents and macaque. Cell Reports 44, 115730 (2025).

50. Mich, J. K., et al. Enhancer-AAVs allow genetic access to oligodendrocytes and diverse populations of astrocytes across species. Preprint at 10.7554/eLife.108640.1 (2025).

51. Mich, J. K. et al. Interneuron-specific dual-AAV *SCN1A* gene replacement corrects epileptic phenotypes in mouse models of Dravet syndrome. Sci. Transl. Med. 17, eadn5603 (2025).

52. Lee, H. et al. Brn3b regulates the formation of fear-related midbrain circuits and defensive responses to visual threat. PLoS Biol 21, e3002386 (2023).

53. Xu, D. et al. SV40 intron, a potent strong intron element that effectively increases transgene expression in transfected Chinese hamster ovary cells. J Cellular Molecular Medi 22, 2231–2239 (2018).

54. Chan, K. Y. et al. Engineered AAVs for efficient noninvasive gene delivery to the central and peripheral nervous systems. Nat Neurosci 20, 1172–1179 (2017).

55. Ivkovic, S. & Ehrlich, M. E. Expression of the Striatal DARPP-32/ARPP-21 Phenotype in GABAergic Neurons Requires Neurotrophins *In Vivo* and *In Vitro*. J. Neurosci. 19, 5409–5419 (1999).

56. Bucci, D. et al. Systematic Morphometry of Catecholamine Nuclei in the Brainstem. Front. Neuroanat. 11, 98 (2017).

57. Fu, Y., Paxinos, G., Watson, C. & Halliday, G. M. The substantia nigra and ventral tegmental dopaminergic neurons from development to degeneration. Journal of Chemical Neuroanatomy 76, 98–107 (2016).

58. Alvarez-Fischer, D. et al. Characterization of the striatal 6-OHDA model of Parkinson’s disease in wild type and α-synuclein-deleted mice. Experimental Neurology 210, 182–193 (2008).

59. Stott, S. R. W. & Barker, R. A. Time course of dopamine neuron loss and glial response in the 6-OHDA striatal mouse model of P arkinson’s disease. Eur J of Neuroscience 39, 1042–1056 (2014).

60. Mendes-Pinheiro, B. et al. Unilateral Intrastriatal 6-Hydroxydopamine Lesion in Mice: A Closer Look into Non-Motor Phenotype and Glial Response. IJMS 22, 11530 (2021).

61. Iancu, R., Mohapel, P., Brundin, P. & Paul, G. Behavioral characterization of a unilateral 6-OHDA-lesion model of Parkinson’s disease in mice. Behavioural Brain Research 162, 1–10 (2005).

62. Yu, C. et al. Striatal mechanisms of turning behaviour following unilateral dopamine depletion in mice. Eur J of Neuroscience 56, 4529–4545 (2022).

63. Ungerstedt, U. Postsynaptic Supersensitivity after 6-Hydroxy-dopamine Induced Degeneration of the Nigro-striatal Dopamine System. Acta Physiologica Scandinavica 82, 69–93 (1971).

64. González-Rodríguez, P. et al. Disruption of mitochondrial complex I induces progressive parkinsonism. Nature 599, 650–656 (2021).

65. Wirthlin, M. E. et al. A Cross-Species Enhancer-AAV Toolkit for Cell Type-Specific Targeting Across the Basal Ganglia. Preprint at 10.64898/2026.02.23.706695 (2026).

66. Ichikawa, T., Ajiki, K., Matsuura, J. & Misawa, H. Localization of two cholinergic markers, choline acetyltransferase and vesicular acetylcholine transporter in the central nervous system of the rat: in situ hybridization histochemistry and immunohistochemistry. Journal of Chemical Neuroanatomy 13, 23–39 (1997).

67. Hol, E. M. & Pekny, M. Glial fibrillary acidic protein (GFAP) and the astrocyte intermediate filament system in diseases of the central nervous system. Current Opinion in Cell Biology 32, 121–130 (2015).

68. Ciesielska, A. et al. Anterograde Axonal Transport of AAV2-GDNF in Rat Basal Ganglia. Molecular Therapy 19, 922–927 (2011).

69. Surdyka, M. M. & Figiel, M. Retrograde capabilities of adeno-associated virus vectors in the central nervous system. BioTechnologia 102, 473–478 (2021).

70. Turcano, P. et al. Levodopa-induced dyskinesia in Parkinson disease: A population-based cohort study. Neurology 91, (2018).

71. Santos-García, D. et al. Levodopa-Induced Dyskinesias are Frequent and Impact Quality of Life in Parkinson’s Disease: A 5-Year Follow-Up Study. Movement Disord Clin Pract 11, 830–849 (2024).

72. Nyholm, D. et al. Pharmacokinetics of Levodopa, Carbidopa, and 3-O-Methyldopa Following 16-hour Jejunal Infusion of Levodopa-Carbidopa Intestinal Gel in Advanced Parkinson’s Disease Patients. AAPS J 15, 316–323 (2013).

73. Bravi, D. et al. Wearing-off fluctuations in Parkinson’s disease: Contribution of postsynaptic mechanisms. Annals of Neurology 36, 27–31 (1994).

74. Picconi, B. et al. Loss of bidirectional striatal synaptic plasticity in L-DOPA–induced dyskinesia. Nat Neurosci 6, 501–506 (2003).

75. Jenner, P. Molecular mechanisms of L-DOPA-induced dyskinesia. Nat Rev Neurosci 9, 665–677 (2008).

76. Muramatsu, S.-I. et al. Behavioral Recovery in a Primate Model of Parkinson’s Disease by Triple Transduction of Striatal Cells with Adeno-Associated Viral Vectors Expressing Dopamine-Synthesizing Enzymes. Human Gene Therapy 13, 345–354 (2002).

77. Samaranch, L. et al. Advancing AADC deficiency therapy through MR-guided multisite delivery of AAV2-hAADC to dopaminergic and serotonergic pathways in the brainstem. Molecular Therapy Advances 34, 201670 (2026).

78. Concha-Marambio, L., Pritzkow, S., Shahnawaz, M., Farris, C. M. & Soto, C. Seed amplification assay for the detection of pathologic alpha-synuclein aggregates in cerebrospinal fluid. Nat Protoc 18, 1179–1196 (2023).

79. Wagner, S. K. et al. Retinal Optical Coherence Tomography Features Associated With Incident and Prevalent Parkinson Disease. Neurology 101, (2023).

80. Svetel, M. et al. Retinal Thickness in Patients with Parkinson’s Disease and Dopa Responsive Dystonia—Is There Any Difference? Biomedicines 13, 1227 (2025).

81. Lim, S. A. O., Kang, U. J. & McGehee, D. S. Striatal cholinergic interneuron regulation and circuit effects. Front. Synaptic Neurosci. 6, (2014).

82. McKinley, J. W. et al. Dopamine Deficiency Reduces Striatal Cholinergic Interneuron Function in Models of Parkinson’s Disease. Neuron 103, 1056–1072.e6 (2019).

83. Ztaou, S. et al. Involvement of Striatal Cholinergic Interneurons and M1 and M4 Muscarinic Receptors in Motor Symptoms of Parkinson’s Disease. Journal of Neuroscience 36, 9161–9172 (2016).

84. Rustay, N. R., Wahlsten, D. & Crabbe, J. C. Influence of task parameters on rotarod performance and sensitivity to ethanol in mice. Behavioural Brain Research 141, 237–249 (2003).

85. Shan, H.-M., Maurer, M. A. & Schwab, M. E. Four-parameter analysis in modified Rotarod test for detecting minor motor deficits in mice. BMC Biol 21, 177 (2023).

86. Tanaka, S., Young, J. W., Halberstadt, A. L., Masten, V. L. & Geyer, M. A. Four factors underlying mouse behavior in an open field. Behavioural Brain Research 233, 55–61 (2012).

87. Liu, Y. et al. Alteration of the cholinergic system and motor deficits in cholinergic neuron-specific Dyt1 knockout mice. Neurobiology of Disease 154, 105342 (2021).

88. Huang, M. et al. Astrocytes in the medial entorhinal cortex are required for spatial exploration. Cell Reports 44, 116173 (2025).

89. Evans, W. R. et al. Functional activation of dorsal striatum astrocytes improves movement deficits in hemi-parkinsonian mice. Preprint at 10.1101/2024.04.02.587694 (2024).

90. Yin, H. H. et al. Dynamic reorganization of striatal circuits during the acquisition and consolidation of a skill. Nat Neurosci 12, 333–341 (2009).

91. Klaus, A., Alves Da Silva, J. & Costa, R. M. What, If, and When to Move: Basal Ganglia Circuits and Self-Paced Action Initiation. Annu. Rev. Neurosci. 42, 459–483 (2019).

92. Alegre-Cortés, J., Sáez, M., Montanari, R. & Reig, R. Medium spiny neurons activity reveals the discrete segregation of mouse dorsal striatum. eLife 10, e60580 (2021).

93. Tranchevent, L.-C., Halder, R. & Glaab, E. Systems level analysis of sex-dependent gene expression changes in Parkinson’s disease. npj Parkinsons Dis. 9, 8 (2023).

94. Schaffner, S. L., Tosefsky, K. N., Inskter, A. M., Appel-Cresswell, S. & Schulze-Hentrich, J. M. Sex and gender differences in the molecular etiology of Parkinson’s disease: considerations for study design and data analysis. Biol Sex Differ 16, 7 (2025).

95. Cortese, M., Riise, T., Engeland, A., Ascherio, A. & Bjørnevik, K. Urate and the risk of Parkinson’s disease in men and women. Parkinsonism & Related Disorders 52, 76–82 (2018).

96. Cerri, S., Mus, L. & Blandini, F. Parkinson’s Disease in Women and Men: What’s the Difference? Journal of Parkinson’s Disease 9, 501–515 (2019).

97. Kent, A. P., Stern, G. M. & Webster, R. A. The effect of benserazide on the peripheral and central distribution and metabolism of levodopa after acute and chronic administration in the rat. British J Pharmacology 100, 743–748 (1990).

98. Florio, E. et al. D2R signaling in striatal spiny neurons modulates L-DOPA induced dyskinesia. iScience 25, 105263 (2022).

99. Francardo, V. et al. Impact of the lesion procedure on the profiles of motor impairment and molecular responsiveness to L-DOPA in the 6-hydroxydopamine mouse model of Parkinson’s disease. Neurobiology of Disease 42, 327–340 (2011).

100. Sebastianutto, I., Maslava, N., Hopkins, C. R. & Cenci, M. A. Validation of an improved scale for rating l-DOPA-induced dyskinesia in the mouse and effects of specific dopamine receptor antagonists. Neurobiology of Disease 96, 156–170 (2016).

101. Daigle, T. L., Wetsel, W. C. & Caron, M. G. Opposite function of dopamine D1 and *N* - methyl-D-aspartate receptors in striatal cannabinoid-mediated signaling. Eur J of Neuroscience 34, 1378–1389 (2011).

