## Supplementary information for "Cell type-selective AADC expression reduces motor deficits and dyskinesia in Parkinson’s disease mouse models" (from another version of this preprint: version 1, 2026-01-21)

Supplemental information

### Enhancer-based AAV approach for selective AADC delivery reduces motor symptoms and dyskinesia in Parkinson's mouse models

Ananya Chowdhury, Alex Fraser, Max Departee, Naz Taskin, Meagan A. Quinlan, John K. Mich, Victoria Omstead, Nathaly Lerma, Ximena Opitz-Araya, Alex C. Hughes, Emily Kussick, Refugio Martinez, Melissa Reding, Elizabeth Liang, Lyudmila Shulga, Dean Rette, Cindy Huang, Brittny Casian, Madison Leibly, Olivia Helback, Tyler Barcelli, Toren Wood, Nicholas Uribe, Colin Bacon, Jessica Bowlus, Dakota Newman, Rana Kutsal, Shannon Khem, Nicholas Donadio, Shenqin Yao, Kara Ronellenfitch, Vonn Wright, Kathryn Gudsnuk, Greg D. Horwitz, Boaz P. Levi, Ed S. Lein Jonathan T. Ting, Tanya L. Daigle

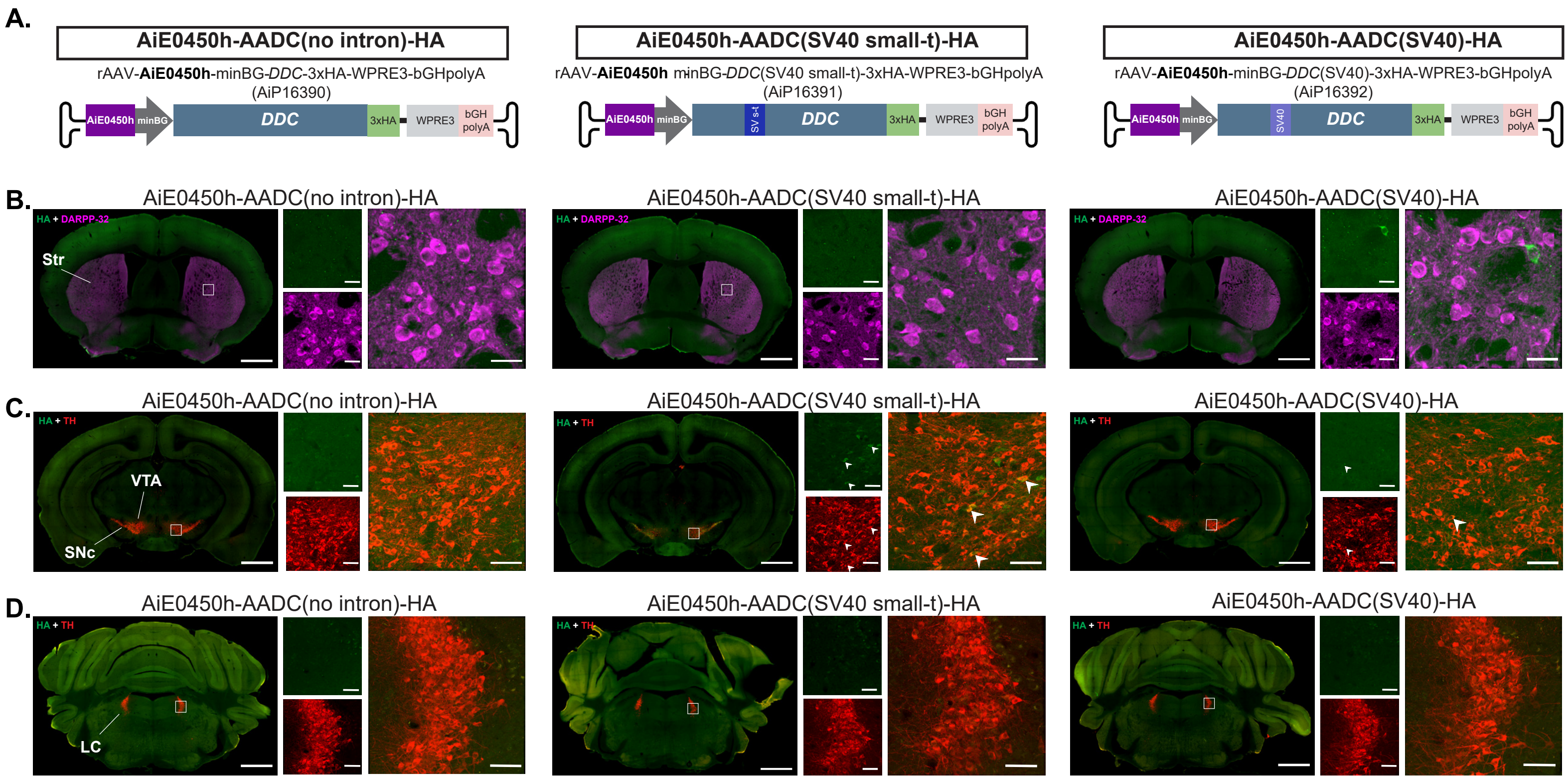


Figure S1: Development and characterization of different AADC transgene designs
(A) Schematic showing the vector designs for AiE0450h-AADC-HA with no intron, SV40 small-t intron, or longer SV40 intronic insertion.
(B) Representative striatal (AP = +0.14 to +0.26 from bregma) coronal images of AADC-HA (green) and DARPP-32+ MSNs (magenta). (C) Representative midbrain (AP = -2.92 to -3.08 from bregma) and (D) representative LC (AP= -5.40 to -5.52 from bregma) coronal images of AADC-HA (green) and TH (red), for AiE0450h-AADC(no intron)-HA (left), AiE0450h-AADC(SV40 small-t)-HA (middle), and AiE0450h-AADC(SV40)-HA (right), following RO administration. Low magnification (10X) stitched, scale bars, 500 μm; High magnification (20X), scale bars, 10 μm for the striatum and 20 μm for VTA and LC.
White arrows depict representative colocalized cells.

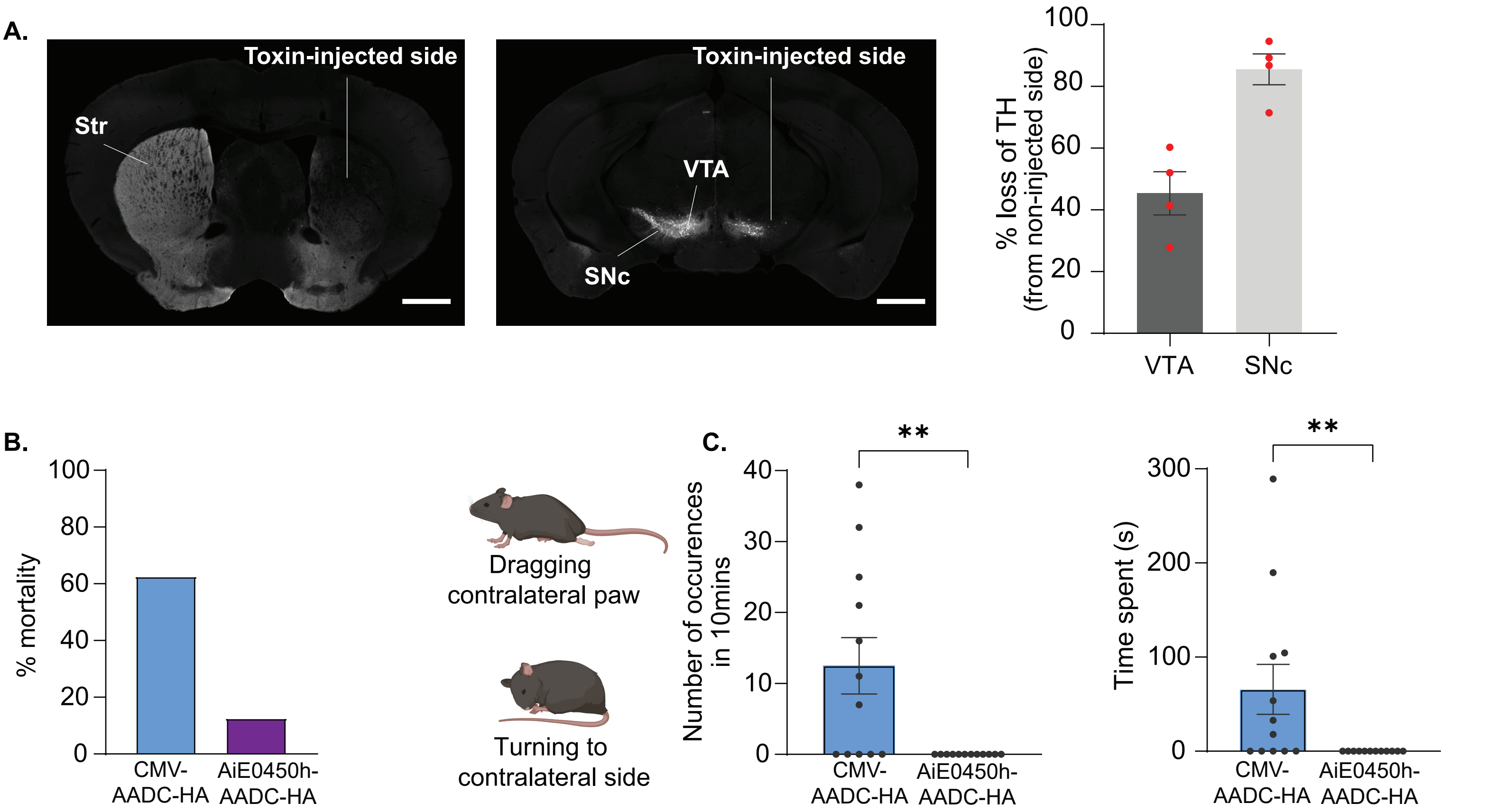


Figure S2: 6-OHDA model validation and phenotypic effect of AADC-AAVs
(A) Representative coronal images of the striatum and midbrain demonstrating the unilateral depletion of TH staining (gray) after toxin administration. Low magnification (10X) stitched, scale bars, 500 µm. Quantification of the loss of TH+ cells in VTA and SNc (% reduction compared to the non-injected hemisphere). Data represented as mean ± SEM. n = 4 mice, each data point average of 3 sections.
(B) Mortality rate 6 weeks after retroorbital injection of the viral vectors (% out of total injected).
(C) Quantification of contralateral asymmetric movements in a cylinder over 10 mins. Data represented as mean ± SEM. n = 12 per group. The p values were calculated by Mann-Whitney test, normality tested with Shapiro-Wilk test. **$p < 0.01$.

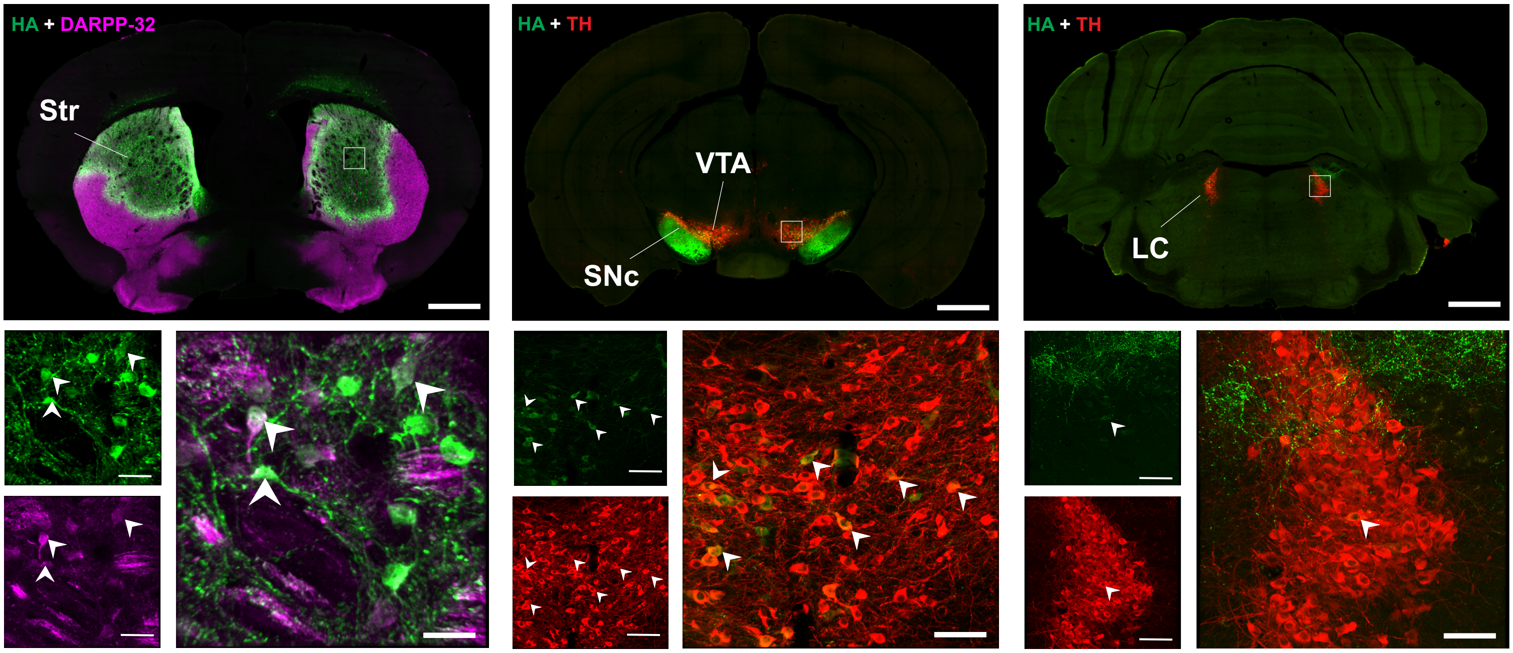


Figure S3: Intra-striatal administration of AiE0450h-AADC-HA at a higher titer (1e10 vg)
Representative coronal images of striatum (AP = +0.14 from bregma) for AiE0450h-AADC-HA showing AADC-HA (green) and DARPP-32+ MSNs (magenta) (left panel), midbrain (AP = -3.08 from bregma) and LC (AP = -5.40 from bregma) showing AADC-HA (green) and TH+ cells (red), following intra-striatal administration of the vector at higher titer of 1e10 vg bilaterally. Low magnification (10X) stitched, scale bars, 500 μm, top; High magnification (20X), scale bars, 10 μm for striatum (bottom left) and 20 μm for midbrain (bottom middle) and LC (bottom right). White arrows depict representative colocalized cells.

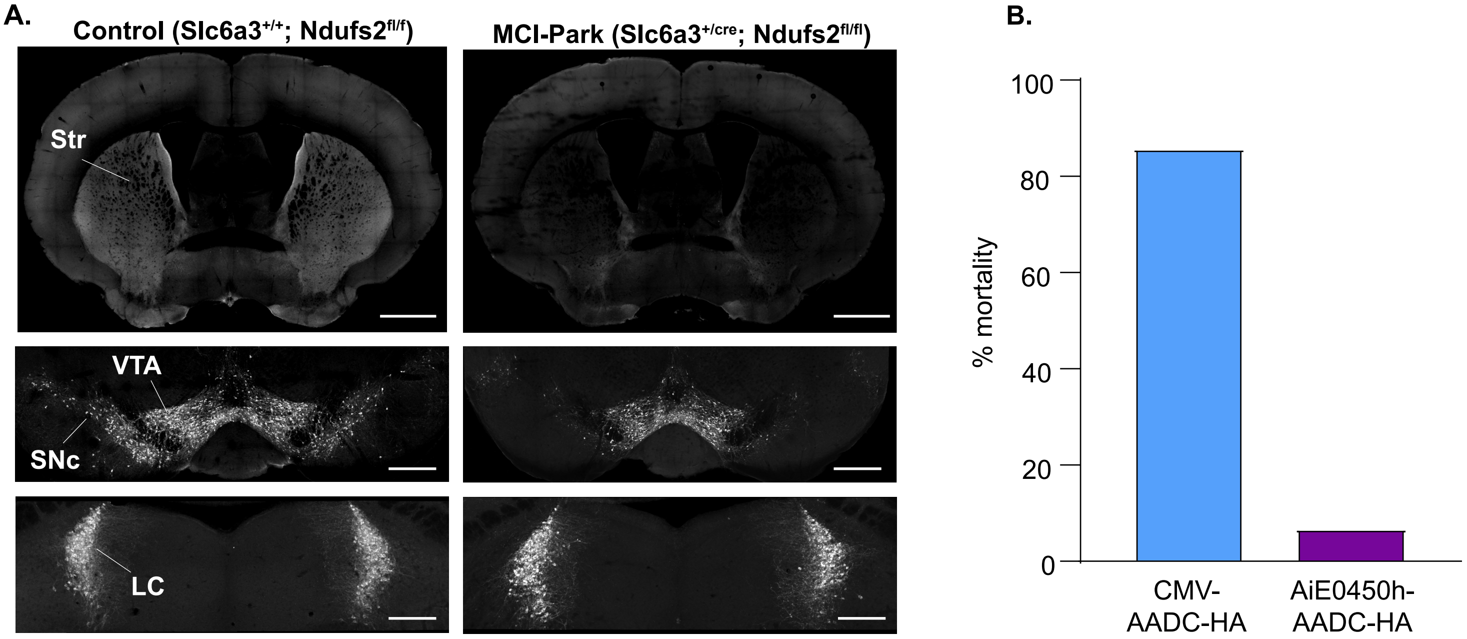


Figure S4: Validation of MCI-Park genetic model and phenotypic observation

(A) Representative coronal images of the striatum (top) and midbrain (bottom) demonstrating the bilateral depletion of TH staining (gray) in MCI-Park ($Slc6a3^{+/cre}$; $Ndufs2^{fl/fl}$, right) compared to the age-matched littermate controls ($Slc6a3^{+/+}$; $Ndufs2^{fl/fl}$, left). Low magnification (10X) stitched, scale bars, 500 μm.

(B) Mortality rate 4 weeks after retroorbital injection of the viral vectors (% out of total injected).

**A.**

| Route | Vector | Completeness VTA | Completeness SNc | Completeness LC | Specificity VTA | Specificity SNc | Specificity LC |
|---|---|---|---|---|---|---|---|
| RO | AiE0450h-AADC-HA | 63.33 | 81.86 | 77.71 | 67.08 | 75.53 | 63.74 |
| RO | CMV-AADC-HA | 41.07 | 67.49 | 41.81 | 73.69 | 57.96 | 48.13 |
| RO | AiE0743m_3xC2-AADC-HA | 37.38 | 72.62 | 47.94 | 43.24 | 57.29 | 63.32 |
| RO | AiE0888m-AADC-HA | 61.06 | 60.33 | 67.17 | 84.35 | 83.45 | 62.07 |
| RO | AiE0387m_3xC1-AADC-HA | 31.29 | 53.21 | 20.36 | 58.20 | 57.27 | 31.08 |
| IS | AiE0450h-AADC-HA | 2.19 | 0.83 | 0.76 | 33.31 | 22.22 | 40.74 |
| IS | AiE0450h-AADC-HA (high) | 18.51 | 27.04 | 6.04 | 50.71 | 69.16 | 56.81 |
| IS | CMV-AADC-HA | 8.80 | 9.04 | 1.53 | 58.09 | 70.42 | 43.80 |

Scale: 0 – 20 – 40 – 60 – 80 – 100

**B.**

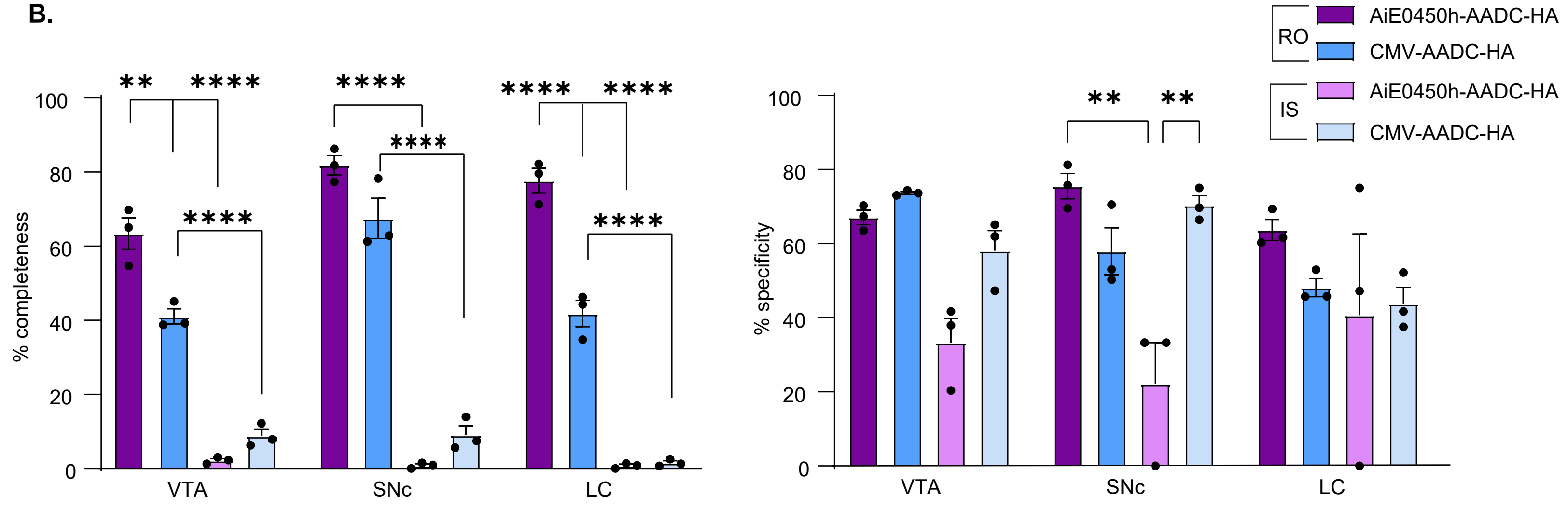


**C.**

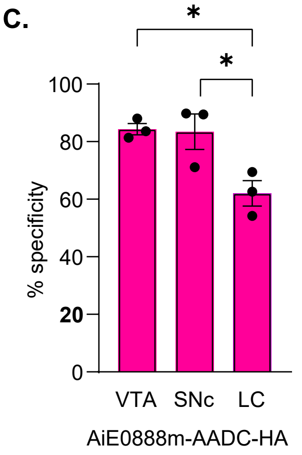


Figure S5: Quantification of AADC-HA colocalization with TH+ neurons in VTA, SNc, and LC for different vectors with the RO and IS routes of administration

(A) Summary heatmap of completeness (left) and specificity (right) for different vectors, determined by IHC with AADC-HA colocalized to the cell-type marker TH+ for catecholaminergic neurons in VTA, SNc, and LC, following the two routes of administration as mentioned. Legend with scale denotes values ranging from 0 (cyan) to 100% (magenta). Percentage values are the average mean value from 3 sections of n = 3 mice each analyzed for all conditions besides AiE0387m_3xC1-AADC-HA (3 sections of n = 2 mice each) and AiE0450h-AADC-HA IS (high) (3 sections of n=1 mouse).

(B) Comparison of % completeness (left) and specificity (right) for AADC-HA colocalized to the cell-type marker TH+ for catecholaminergic neurons in VTA, SNc, and LC, following the two routes of administration with AiE0450h-AADC-HA and CMV-AADC-HA. Data represented as mean ± SEM, n = 3 mice, each data point average of 3 sections. The p values were calculated by two-way ANOVA followed by Sidak's post hoc test. $^{**}p < 0.01$, $^{****}p < 0.0001$.

(C) Comparison of % specificity for AADC-HA colocalized to the cell-type marker TH+ for catecholaminergic neurons in VTA, SNc, and LC, following RO administration with AiE0888m-AADC-HA. Data represented as mean ± SEM, n = 3 mice, each data point average of 3 sections. The p values were calculated by one-way ANOVA followed by Sidak's post hoc test. $^{*}p < 0.05$.

**Table S1: Summary table of behavioral results and mortality rates observed in 6-OHDA and MCI-Park models with AAVs delivered via two routes of administration.** Rescued = statistically significant reversal of behavioral deficit with AAV treatment. No rescue = no change in the behavioral deficit with AAV treatment. No asymmetry = AAV did not produce this behavioral effect. Increased = AAV produced notable rotations. NA = data is not available and N/A = not applicable for the model

| PD model | Route of admin | Vector | Target cell types | Rotarod | Open Field | Contralateral paw touches | Contralateral rotations | LID | Mortality % (above baseline) |
|---|---|---|---|---|---|---|---|---|---|
| 6-OHDA | RO | AiE450h-SYFP2 | MSN | No rescue | No rescue | No rescue | NA | NA | 0 |
| | | AiE450h-AADC-HA | MSN | Rescued | No rescue | Rescued | No asymmetry | NA | 5-20 |
| | | CMV-AADC-HA | Ubiquitous | No rescue | No rescue | NA | Increased | NA | >50 |
| | | AiE0743m-AADC-HA | Striatal cholinergic | Rescued | No rescue | NA | NA | NA | 0 |
| | | AiE0888m-AADC-HA | Midbrain dopaminergic | Rescued | Rescued | NA | NA | NA | 0 |
| | | AiE0387m-AADC-HA | Astrocytes | No rescue | Rescued | NA | NA | NA | 0 |
| | IS | AiE0450h-AADC-HA | MSN | Rescued | No rescue | Rescued | NA | Rescued | 0 |
| | | CMV-AADC-HA | MSN | Rescued | No rescue | Rescued | NA | No rescue | 0 |
| MCI-Park | RO | AiE0450h-AADC-HA | MSN | Rescued | Rescued | N/A | N/A | NA | 5-20 |
| | | AiE0743m-AADC-HA | Striatal cholinergic | No rescue | No rescue | N/A | N/A | NA | 5-20 |
| | | AiE0888m-AADC-HA | Midbrain dopaminergic | No rescue | No rescue | N/A | N/A | NA | 5-20 |
| | | CMV-AADC-HA | Ubiquitous | Small n | | | | | >80 |
| | IS | AiE0450h-AADC-HA | MSN | Rescued | Rescued | N/A | N/A | NA | 5-20 |
| | | CMV-AADC-HA | Ubiquitous | Rescued | No rescue | N/A | N/A | NA | 5-20 |

**Table S2: Reagents and resources used in this study**

| REAGENT or RESOURCE | SOURCE | IDENTIFIER |
|---|---|---|
| Antibodies | | |
| Chicken Anti-Tyrosine Hydroxylase antibody | Abcam | AB76442, RRID:AB_10412418 |
| HA-Tag (C29F4) Rabbit mAb | Cell Signaling Technology | 3724, RRID:AB_1549585 |
| BD Transduction Laboratories™ Purified Mouse Anti-DARPP-32 | BD Biosciences | 611520, RRID:AB_398980 |
| Anti-CHAT Antibody Mouse IgG2B Clone CL3173 | Atlas Antibodies | AMAb91130, RRID:AB_2665812 |
| Anti-Glial Fibrillary Acidic Protein Antibody, clone GA5 in Mouse | MilliporeSigma | MAB360, RRID:AB_3695585 |
| Anti-Green Fluorescent Protein Antibody, Chicken Polyclonal | Aves | GFP-1020, RRID:AB_10000240 |
| Rabbit Anti-NeuN Polyclonal Antibody | MilliporeSigma | ABN78, RRID:AB_10807945 |
| Goat α-Rabbit IgG AF488 | Thermo Fisher Scientific | A11008, RRID:AB_143165 |
| Goat α-Mouse IgG12B AF555 | Thermo Fisher Scientific | A21147, RRID:AB_2535783 |
| Goat α-Chicken IgY AF488 | Thermo Fisher Scientific | A11039, RRID:AB_2534096 |
| Goat α-Chicken IgY AF568 | Thermo Fisher Scientific | A11041, RRID:AB_2534098 |
| Goat α-Rabbit IgG AF568 | Thermo Fisher Scientific | A11011, RRID:AB_2534078 |
| Goat α-Mouse IgG1 AF647 | Thermo Fisher Scientific | A21240, RRID:AB_2535809 |
| Anti-HA.11 Epitope Tag Mouse Monoclonal | Biolegend | 901513, RRID:AB_2565335 |
| DDC (D6N8N) Rabbit mAb | Cell Signaling Technology | 13561S, RRID:AB_2798255 |
| Anti-Actin Antibody, clone C4 | MilliporeSigma | MAB1501R, RRID:AB_2223041 |
| Purified Mouse Anti-DARPP-32 | BD Biosciences | 611520, RRID:AB_398980 |
| IRDYE® 680RD Donkey anti-mouse IGG secondary antibody | LICORbio | 926-68072, RRID:AB_10953628 |
| IRDye® 800CW Donkey anti-Rabbit IgG secondary Antibody | LICORbio | 926-32213, RRID:AB_621848 |
| Bacterial and virus strains | | |
| One Shot™ Stbl3™ chemically competent *E. coli* | Thermo Fisher Scientific | C737303 |
| Chemicals, peptides, and recombinant proteins | | |
| 1x PBS | Corning | 21-040-CV |
| Sodium chloride solution (0.9%), BioXtra, suitable for cell culture | Sigma | S8776 |

| | | |
|---|---|---|
| Sodium azide 5% | Ricca Chemical Company | 7144.8-16 |
| 4',6-diamidino-2-phenylindole (DAPI) | Fisher | 62248 |
| Bovine Serum Albumin | Sigma | A3294 |
| Normal Goat Serum | Vector Laboratories | S-1000 |
| TritonX100 | VWR | 97062-208 |
| 6-Hydroxydopamine hydrochloride | Sigma | H4381 |
| L-Ascorbic Acid | Sigma | A7506 |
| Lactated Ringers Solution | Vedco | 50989-883-17 |
| Rimadyl (carprofen) | Zoetis | 151-199 |
| Levodopa | Millipore Sigma | PHR1271 |
| Benserazide Hydrochloride | Millipore Sigma | B7283 |
| 4x Laemmeli buffer with 5% BME | Thermo Fisher Scientific | J60015.AD |
| Novex 10% Tris/Glycine gels | Thermo Fisher Scientific | XP00100BOX |
| Intercept® (TBS) Blocking Buffer | LICORbio | 927-60001 |
| Nitrocellulose membranes | Thermo Fisher Scientific | 88025 |
| Critical commercial assays | | |
| BCA protein assay | Thermo Fisher Scientific | 23225 |
| HiSpeed Plasmid Maxi Kit | Qiagen | 12663 |
| Experimental models: Organisms/strains | | |
| C57BL/6J mice | The Jackson Laboratory | IMSR_JAX:000664 |
| B6.SJL-Slc6a3tm1.1(cre)Bkmn/J | The Jackson Laboratory | IMSR_JAX:006660 |
| Ndufs2tm1.1Job Slc6a3tm1.1(wt)Bkmn/SurmJ | The Jackson Laboratory | IMSR_JAX:036313 |
| Macaca nemestrina animals | Washington National Primate Research Center | N/A |
| Recombinant DNA | | |
| AiP12236 - pAAV-AiE0450h-minβG-SYFP2-WPRE3-BGHpA (Alias: CN2236) | Addgene; Hunker et.al[40] | 224160 |
| AiP13663 - pAAV-AiE0450h-minβG-DDC(HBB3)-3xHA-WPRE3-bGHpA (Alias: CN3663) | Addgene; this study | In progress |
| AiP16328 - pAAV-CMV-DDC(HBB3)-3xHA-WPRE3-bGHpA (Alias: HCT104) | Addgene; this study | In progress |
| AiP13912 - pAAV-AiE0743m_3xCore2-minßG-DDC(HBB3)-3xHA-WPRE3-bGHpA (Alias: CN3912) | Addgene; this study | In progress |
| AiP13742 - pAAV-AiE0888m-minßG-DDC(HBB3)-3xHA-WPRE3-bGHpA (Alias: CN3742) | Addgene; this study | In progress |

| | | |
|---|---|---|
| AiP15007 - pAAV-AiE0387m_3xCore1-minßG-DDC(HBB3)-3xHA-WPRE3-bGHpolyA (Alias: CN5007) | Addgene; this study | In progress |
| AiP16390 - pAAV-AiE0450h-minβG-DDC-3xHA-WPRE3-bGHpA (Alias: HCT102) | Addgene; this study | In progress |
| AiP16391 - pAAV-AiE0450h-minβG-DDC(GENSAT)-3xHA-WPRE3-bGHpA (Alias: HCT120) | Addgene; this study | In progress |
| AiP16392 - pAAV-AiE0450h-minβG-DDC(SV40)-3xHA-WPRE3-bGHpA (Alias: HCT121) | Addgene; this study | In progress |
| Software and algorithms | | |
| Prism 10.4.1 | Dotmatics | SCR_002798 |
| Imaris 10.0.1 | Bitplane AG | SCR_007370 |
| BioRender | Biorender | SCR_018361 |
| Ethovision | Noldus | SCR_000441 |
| ImageJ | National Institutes of Health | SCR_003070 |
| Other | | |
| Rotarod | Med Associates | ENV-574M |
| Open Field | Maze Engineers/Noldus | Multi-arena open field |
| Cylinder | N/A | NA |
| Model 1900 Stereotaxic Alignment Instrument | David Kopf Instruments | 1900 |
| Nanoject II | Drummond, now discontinued | 3-000-204 |
| Nanoject Controller | World Precision Instrument | SYS-MICRO4 |
| Freezing microtome | Leica | SM2000R/SM2010R |
| Brainsight Injection Robot | Rogue Research | Vet Robot |
| Prolong Gold Antifade Mounting Medium | Invitrogen | P36930 |
| Tissue-Tek O.C.T Compound | Sakura | 4583 |
| D12492i Rodent Diet with 60 kcal% Fat | Research Diets | D12492i |
| Mineral oil | Sigma | M8410 |
| Betadine Solution | Avrio Health | 67618-150-17 |
| PROLENE® Polypropylene Suture | Ethicon | 8661G |
| Isoflurane | Pivetal | 46066-115-04 |
| DietGel Boost | ClearH20 | 72-04-5022 |
| HydroGel | ClearH20 | 70-01-1062 |
| Proparacaine Hydrochloride Ophthalmic Solution 0.5% | Bausch + Lomb | 24208-730-06 |
